# Blocking palmitoylation of *Toxoplasma gondii* myosin light chain 1 disrupts glideosome composition but has little impact on parasite motility

**DOI:** 10.1101/2020.08.13.250399

**Authors:** Pramod K. Rompikuntal, Ian T. Foe, Bin Deng, Matthew Bogyo, Gary E. Ward

## Abstract

*Toxoplasma gondii* is a widespread apicomplexan parasite that causes severe disease in immunocompromised individuals and the developing fetus. Like other apicomplexans, *T. gondii* uses an unusual form of gliding motility to invade cells of its hosts and to disseminate throughout the body during infection. It is well established that a myosin-based motor consisting of a Class XIVa heavy chain (TgMyoA) and two light chains (TgMLC1 and TgELC1/2) plays an important role in parasite motility. The ability of the motor to generate force at the parasite periphery is thought to be reliant upon its anchoring and immobilization within a peripheral membrane-bound compartment, the inner membrane complex (IMC). The motor does not insert into the IMC directly; rather, this interaction is believed to be mediated by the binding of TgMLC1 to the IMC-anchored protein, TgGAP45. The binding of TgMLC1 to TgGAP45 is therefore considered a key element in the force transduction machinery of the parasite. TgMLC1 is palmitoylated, and we show here that palmitoylation occurs on two N-terminal cysteine residues, C8 and C11. Mutations that block TgMLC1 palmitoylation disrupt the association of TgMLC1 with the membrane fraction of the parasite in phase partitioning experiments and completely block the binding of TgMLC1 to TgGAP45. Surprisingly, the loss of TgMLC1 binding to TgGAP45 in these mutant parasites has little effect on their ability to initiate or sustain movement. These results question a key tenet of the current model of apicomplexan motility and suggest that our understanding of gliding motility in this important group of human and animal pathogens is not yet complete.

**Importance:** Gliding motility plays a central role in the life cycle of *T. gondii* and other apicomplexan parasites. The myosin motor thought to power motility is essential for virulence but distinctly different from the myosins found in humans. Consequently, an understanding of the mechanism(s) underlying parasite motility and the role played by this unusual myosin may reveal points of vulnerability that can be targeted for disease prevention and treatment. We show here that mutations that uncouple the motor from what is thought to be a key structural component of the motility machinery have little impact on parasite motility. This finding runs counter to predictions of the current, widely-held “linear motor” model of motility, highlighting the need for further studies to fully understand how apicomplexan parasites generate the forces necessary to move into, out of and between cells of the hosts they infect.

## Introduction

Toxoplasmosis is among the most widespread and common parasitic infections of humans (1). Acute infection, while typically subclinical and self-limiting, can cause life-threatening disease in immunocompromised individuals and the developing fetus. The causative agent of toxoplasmosis is the protozoan parasite, *Toxoplasma gondii. T. gondii* and other parasites of the phylum Apicomplexa, including those that cause malaria and cryptosporidiosis, use an unusual form of substrate-dependent gliding motility to invade into and egress from host cells, migrate across biological barriers, and disseminate through the infected host’s tissues (2-4).

Gliding motility in apicomplexan parasites is controlled, at least in part, by an unconventional class XIVa myosin, MyoA (5-15). According to the “linear motor” model of motility that has dominated the field for the last decade (reviewed in (16); see Fig. 1A), *T. gondii* MyoA (TgMyoA) and its associated light chains (TgMLC1 and either TgELC1 or TgELC2) are anchored to the parasite’s inner membrane complex (IMC) via the acylated <u>g</u>lideosome-<u>a</u>ssociated <u>p</u>rotein, TgGAP45. TgGAP45, in turn, binds to the transmembrane proteins TgGAP40 and TgGAP50. TgGAP50 is firmly immobilized within the IMC lipid bilayer, potentially serving as a fixed anchor against which the motor complex can generate force (17, 18). This large, heterooligomeric protein complex (TgMyoA, its light chains, TgGAP40, TgGAP45 and TgGAP50) is referred to as “the glideosome”. In the linear motor model, short actin filaments located between the parasite plasma membrane and the IMC are connected to ligands on the substrate through a glideosome-associated connector protein (GAC; (19)) that binds to the cytosolic tails of surface adhesins. Because the motor is anchored into the IMC, when the motor displaces the fixed actin filaments rearward, the parasite moves forward relative to the substrate (Fig. 1A).

**Figure 1.**
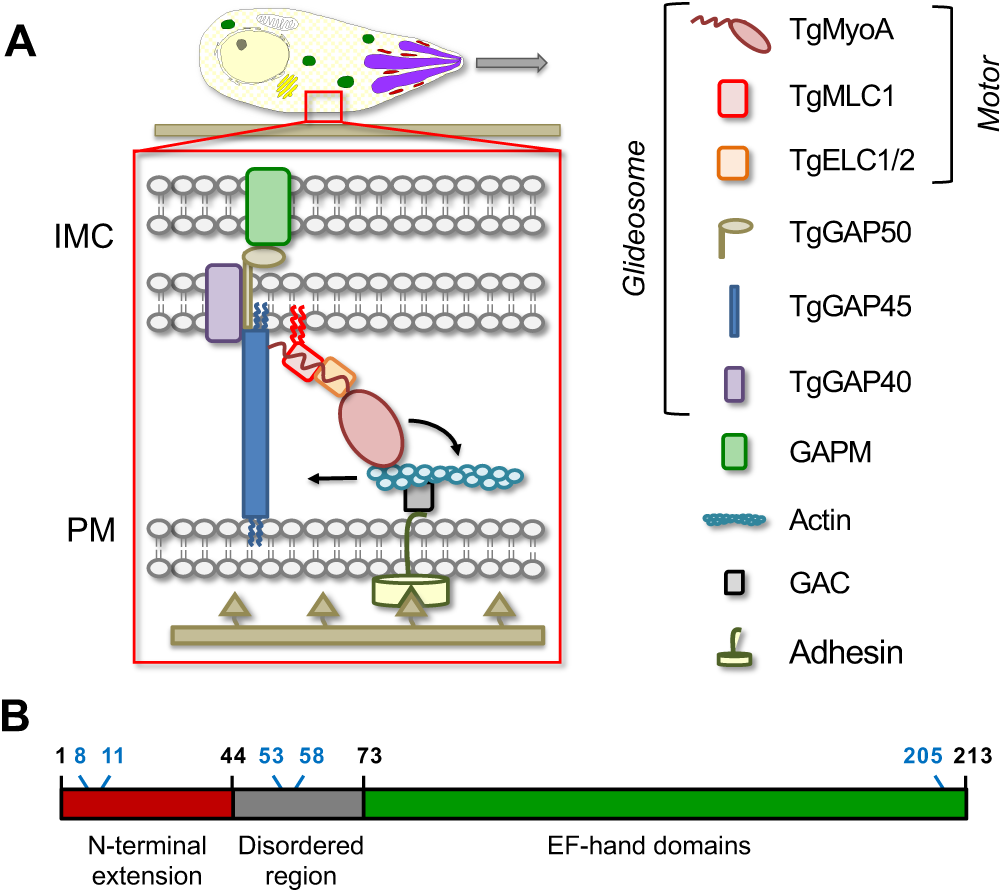
Schematic illustrations of the glideosome and TgMLC1 domain structure. **(A)** In the linear motor model of motility (reviewed in (16)), the TgMyoA motor (TgMyoA and its associated light chains, TgMLC1 and either TgELC1 or TgELC2) is anchored to the parasite’s inner membrane complex (IMC) via the acylated <u>g</u>lideosome-<u>a</u>ssociated <u>p</u>rotein, TgGAP45, and the transmembrane proteins TgGAP40 and TgGAP50. The lumenal portion of GAP50 is thought to interact with GAPM, a protein that spans the inner IMC membrane, and which likely connects the entire glideoeome to the underlying parasite cytoskeleton. Short actin (TgACT1) filaments located between the parasite plasma membrane and the IMC are connected to ligands on the substrate through a linker protein, possibly the <u>g</u>lideosome-<u>a</u>ssociated <u>c</u>onnector protein (GAC), which binds to the cytosolic tails of surface adhesins such as TgMIC2. The TgMyoA motor displaces the actin filaments rearward; because the motor is connected to the IMC and the actin is connected to the substrate, this causes the parasite to move forward relative to the substrate. The depiction of a pair of acyl chains on the N-terminus of TgMLC1 (red squiggles) and their interaction with the IMC membrane is based on results reported here. **(B)** TgMLC1 consists of a 44-amino acid N-terminal extension, a central disordered region, and four EF hand-like domains, which interact with the tail of TgMyoA. The positions of the five cysteines in the protein are shown in blue; CSS-Palm 4.0 predicted C8 and C11 as likely sites of palmitoylation.

TgMLC1 is thought to play two key roles within the *T. gondii* glideosome. First, TgMLC1 binds to the C-terminal tail of TgMyoA to reinforce the motor’s lever arm (10, 13, 20). The lever arm amplifies small motions at the myosin active site into larger movements that are capable of displacing actin filaments (10, 21). Consistent with this proposed function, recombinant TgMyoA is inactive in *in vitro* motility assays in the absence of TgMLC1 ((10) and unpublished data). Second, an interaction between the N-terminal portion of TgMLC1 and the C-terminal portion of TgGAP45 is believed to be the critical link that tethers the motor to the IMC (Fig. 1A; (22, 23). Given these proposed functions, is not surprising that TgMLC1 is an essential protein, and parasites depleted of TgMLC1 are significantly impaired in 3D motility, invasion, and host cell egress (24, 25).

While the importance of TgMyoA, TgMLC1 and the other glideosome components in motility is well established, recent data have called into question whether they are organized and function as described by the linear motor model and/or whether alternative motility mechanisms exist (24-29). For example, the ability of apicomplexan parasites to rock back and forth on a substrate along their anterior to posterior axis (30-36) is hard to reconcile with the linear motor model, as is the ability of parasites engineered to lack key components of the glideosome to continue moving ((24-26); see also (37)). Given the central importance of motility in the parasite’s life cycle and virulence, it is important to fully understand how these proteins work together to generate the forces required to drive parasite movement.

S-palmitoylation is the reversible covalent attachment of a 16-carbon saturated fatty acid via a thioester linkage to cysteine residues of integral and peripheral membrane proteins (38, 39). This widespread post-translational modification of proteins mediates membrane association and can regulate subcellular localization, trafficking, structure, stability and diverse aspects of protein function (38, 40-42). Palmitoylation is thought to play an important role in the biology of *T. gondii* and other apicomplexan parasites (43-54). Recent chemical proteomic studies identified several hundred putatively palmitoylated proteins in *T. gondii* (282 unique proteins in one study (54) and 401 in another (44)). Surprisingly, these proteins included all components of the glideosome, including TgMLC1 (44, 54).

TgMLC1 contains five cysteine residues (Fig. 1B), two of which (C8 and C11) are predicted by CSS-Palm 4.0 to be potential sites of palmitoylation. These two cysteines are found within the apicomplexan-specific N-terminal extension of TgMLC1 (Fig. 1B), which is the region of the protein that binds to TgGAP45 ((23); Fig. 1A). Given the important role that TgMLC1 is thought to play in TgMyoA function and motility, we sought to experimentally confirm C8 and/or C11 as the sites of TgMLC1 palmitoylation and to explore the phenotypic consequences of mutations that block this modification.

## Results

### Identification of the sites of palmitoylation on TgMLC1

To determine whether C8 and/or C11 are sites of palmitoylation on TgMLC1, we replaced the endogenous *TgMLC1* gene with mutant alleles that produce either single (C8S, C11S) or double (C[8,11]S) cysteine to serine mutations, rendering these sites non-palmitoylatable. Each mutant protein was also FLAG-tagged at its N-terminus (Fig. S1; see Table S1 for a list of parasite strains used in this study and their designations). A fourth parasite line expressing FLAG-tagged wild-type TgMLC1 was similarly generated (WT). To determine the effect (if any) of the mutations on TgMLC1 palmitoylation, WT, C8S, C11S and C(8,11)S parasites were grown in the palmitic acid analog, 17-octadecynoic acid (17-ODYA). FLAG-tagged TgMLC1 was then immunoprecipitated and subjected to SDS-PAGE. Because 17-ODYA contains a terminal alkyne group, it can be fluorescently tagged with rhodamine-azide through a copper-catalyzed cycloaddition reaction; the amount of rhodamine bound to proteins in the immunoprecipitate can then be visualized by fluorescence scanning (54). The amount of rhodamine fluorescence associated with TgMLC1 (31 kDa) was significantly reduced in both the C8S and C11S mutants compared to WT, with C8S showing a greater reduction than C11S (Fig. 2). In the C(8,11)S double mutant, no 17-ODYA TgMLC1 labeling above background was detectable. In a previous study, C8 and/or C11 were speculated to be sites of palmitoylation on TgMLC1, and parasites expressing a second copy of TgMLC1 in which these two cysteines were mutated to alanines were generated (23). To compare our results to theirs, we generated a C(8,11)A allelic replacement line and found that, like the C(8,11)S double mutation, the C(8,11)A double mutation completely blocked 17-ODYA labeling (Fig. 2). Taken together, these data identify C8 and C11 as essential for, and very likely the sites of, palmitoylation on TgMLC1.

**Figure 2.**
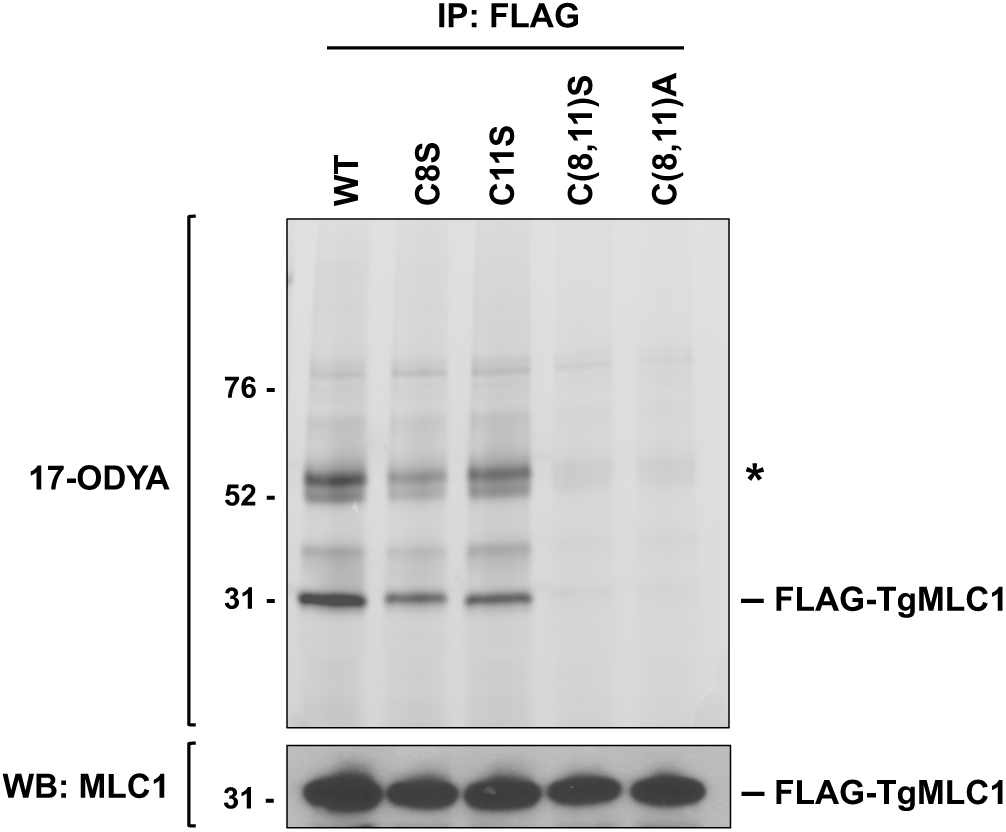
Cys8 and Cys11 are the likely sites of palmitoylation on TgMLC1. Parasites expressing either wild-type (WT) or mutant FLAG-tagged TgMLC1 were labeled with the palmitic acid analog 17-ODYA, and anti-FLAG resin was then used to pull down FLAG-TgMLC1 and associated proteins. The proteins in the pulldown were resolved by SDS-PAGE and visualized either by rhodamine fluorescence scan (upper panel), to show the position of 17-ODYA in the gel, or by western blotting with anti-TgMLC1 (lower panel). Numbers on the left indicate molecular mass in kDa; only the ∼30kDa portion of the western blot is shown. The predominant ∼31-kDa ODYA-labeled band comigrates with FLAG-TgMLC1; its labeling intensity is reduced in the C8S and C11S single mutants and abolished completely in the C(8,11)S and C(8,11)A double mutants. The western blot shows similar protein loads in all samples. The asterisk indicates a doublet of ODYA-labeled proteins at ∼50kDa that is pulled down with wild-type FLAG-TgMLC1 but not with either of the double mutants – see text for details.

### Subcellular localization of non-palmitoylatable TgMLC1

TgMLC1 normally localizes uniformly around the parasite periphery (55). It was previously reported that the C(8,11)A double mutation caused TgMLC1 to mislocalize to the cytosol (23). It was therefore surprising that, in our hands, both the C(8,11)S and C(8,11)A mutant proteins remained localized at the parasite periphery (Fig. 3). Re-examination of the images shown in Frenal *et al*. (2010) revealed that most of the C(8,11)A mutant protein (named MLC1^CC-AA^ in that study) was indeed also found at the parasite periphery, although there was a minor amount in the cytosol (for comparison, see the localization of a different mutant in that same study, MLC1-PGF^AIA^, which was clearly cytosolic (23)). The fact that we detect little to no cytosolic staining with the C(8,11)A allele may reflect differences between the two studies in protein expression levels, since our mutant protein was expressed from the endogenous promoter at the endogenous locus whereas the previous study expressed the mutant gene in parasites also expressing the wild-type allele (23). Taken together, these data show that mutations that block TgMLC1 palmitoylation do not appreciably alter its localization at the parasite periphery.

**Figure 3.**
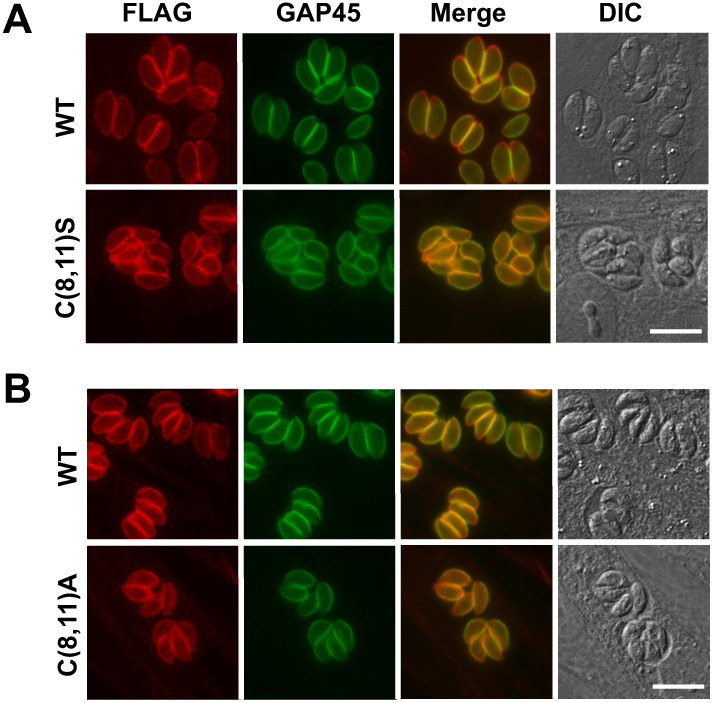
C(8,11)S and C(8,11)A mutations do not affect localization of TgMLC1 to the parasite periphery. Infected HFF cells were fixed, permeabilized and stained for FLAG-tagged TgMLC1 (red) or TgGAP45 (green). The corresponding merged and DIC images are also shown. Upper panels compare the localization of WT *vs*. C(8,11)S TgMLC1; lower panels compare WT *vs*. C(8,11)A TgMLC1. Scale bar = 10 μm.

### Blocking palmitoylation of TgMLC1 alters its phase partitioning in Triton X-114

Next, we tested whether the mutations that block TgMLC1 palmitoylation within the parasite alter the phase partitioning of the protein in the non-ionic detergent Triton X-114 (TX-114). TX-114 efficiently solubilizes most proteins in the parasite at 4°C; when subsequently warmed above the cloud point of the detergent (20°C), intermicellar interactions cause the solution to separate into aqueous and detergent phases, which are enriched in hydrophilic and integral membrane proteins, respectively (56, 57). WT, C8S and C11S TgMLC1 each partition roughly equally into the aqueous and detergent phases, but the C(8,11)S TgMLC1 double mutant is found almost entirely in the aqueous phase (Figs. 4A and S2), suggesting a lack of direct membrane association in the absence of palmitoylation. Similar results were seen with the C(8,11)A double mutant (Fig. 4A). As a control, the same samples were probed for TgGRA8, a dense granule protein (58) unrelated to TgMLC1. As expected, the phase partitioning of TgGRA8 was relatively unaffected by the TgMLC1 mutations (Figs. 4B and S2). These data suggest that the peripheral localization we observed for the C(8,11)S and C(8,11)A TgMLC1 mutants is likely mediated by interaction with other membrane-associated protein(s), rather than by a direct association with the lipid bilayer itself.

**Figure 4.**
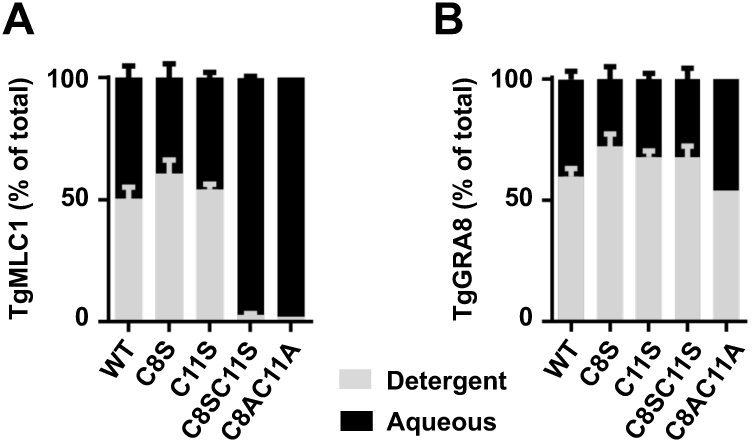
Blocking TgMLC1 palmitoylation causes the protein to shift into the aqueous phase in Triton X-114. WT, C8S, C11S, C(8,11)S and C(8,11)A parasites were extracted at 4°C in Triton X-114 and the extracted proteins phase partitioned by shifting the temperature to 20°C. The amounts of **(A)** TgMLC1and **(B)** TgGRA8 recovered in the detergent (grey) and aqueous (black) phases from each sample were determined by quantitative western blotting (see Fig. S2 for a representative western blot), and are displayed here as the percentage of the total TgMLC1 recovered in the two phases combined. The data shown are the means and standard error of the mean (SEM) from 2 (C8S, C11S) or 4 (WT, C[8,11]S) independent replicates; C(8,11)A parasites were analyzed once.

### Effects of TgMLC1 palmitoylation on the composition of the glideosome

In the 17-ODYA labeling experiments, two prominently labeled ∼50 kDa proteins were recovered in the FLAG-WT pulldowns in addition to FLAG-tagged TgMLC1, and these bands were not present in pulldowns from either the C(8,11)S or C(8,11)A double mutant (Fig. 2, asterisk). It was previously suggested that proteins of this size copurifying with WT TgMLC1 but not C(8,11)A might be other members of the glideosome complex (23). This hypothesis was strengthened by our subsequent observation that most, if not all, glideosome components are palmitoylated (54). We therefore analyzed the FLAG pulldowns of C8S, C11S and C(8,11)S parasites by western blot with antibodies against TgGAP45, TgELC1 and TgMyoA. As expected, TgGAP45 was recovered in the FLAG pulldown from parasites expressing WT TgMLC1; in striking contrast, virtually no TgGAP45 was recovered in FLAG pulldowns from parasites expressing C(8,11)S TgMLC1 (Fig. 5A). Pulldowns from parasites expressing the C8S and C11S single mutations contain intermediate levels of TgGAP45. Quantification of the western blot signals confirmed these observations, and revealed that concomitant with the decrease in TgGAP45 in the IP of the double mutant, there was a 2-3-fold <u>in</u>crease in the amount of TgELC1 and TgMyoA recovered (Fig. 5B). Similar results were seen with the C(8,11)A mutant (Fig. S3). The lack of TgGAP45 in the IP from the double mutant is not due to changes in TgGAP45 expression in this parasite line, as western blots of parasite lysate before immunoprecipitation show similar amounts of TgGAP45 (Fig. 5A and S3, “input”). Similarly, anti-TgMyoA western blots of whole parasite lysates reveals no changes in the level of expression of TgMyoA in either the C(8,11)S or C(8,11)A mutant parasite lines (Fig. S4). Blocking TgMLC1 palmitoylation seems to therefore block its ability to interact with TgGAP45, while simultaneously increasing its interaction with TgMyoA and TgELC1.

**Figure 5.**
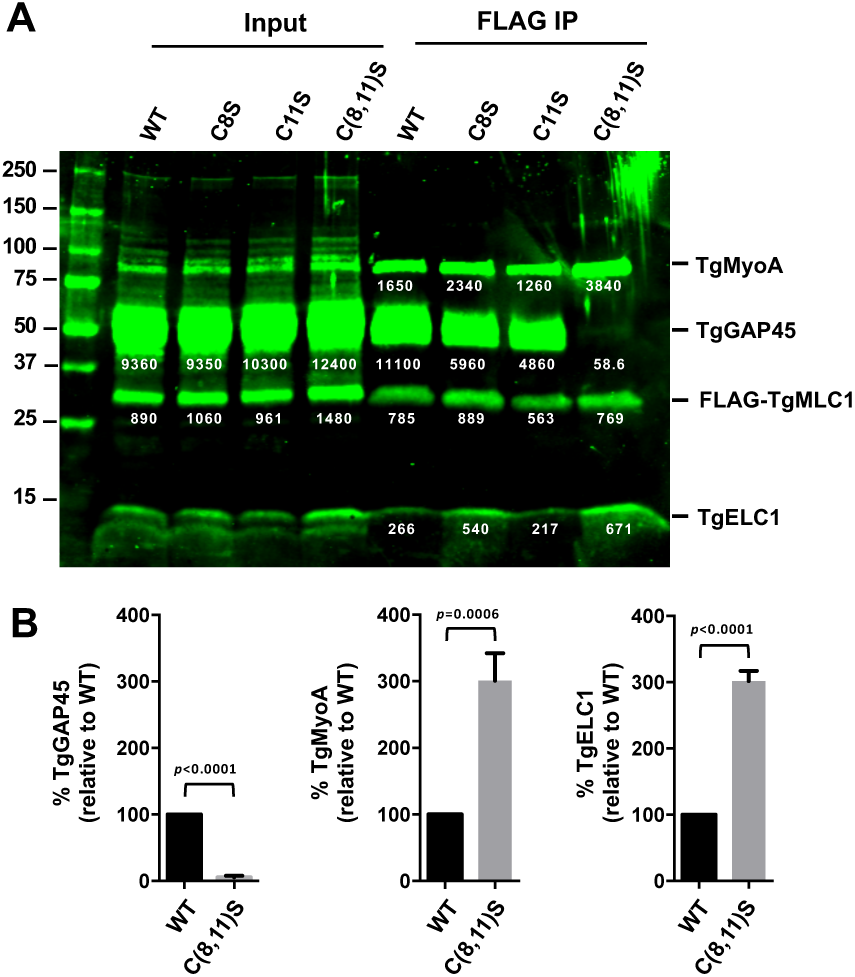
Blocking TgMLC1 palmitoylation alters the composition of the glideosome. **(A)** Parasites expressing wild-type or mutant FLAG-tagged TgMLC1 were gently extracted in Triton X-100, and the soluble proteins (Input) were used for anti-FLAG immunoprecipitations (FLAG IP). Immunoprecipitated proteins were resolved by SDS-PAGE and analyzed by sequential western blotting with anti-TgMyoA, -TgGAP45, -TgMLC1 and -TgELC1; the fluorescent signal intensity of each immunoreactive band is indicated by the white number below the band (relative fluorescence units). Numbers on the left indicate molecular mass in kDa. **(B)** The signal intensities of TgGAP45, TgMyoA and TgELC1 immunoprecipitated from the C(8,11)S mutant parasite line is shown relative the corresponding band from the WT line, after normalizing to the amount of TgMLC1 recovered in each sample. Shown are the means and SEM from six independent experiments; differences were assessed using an unpaired two-tailed t-test.

### Effect of TgMLC1 palmitoylation on parasite motility

Given the dramatic effect of the C(8,11)S double mutation on the binding of TgMLC1 to TgGAP45, we expected to see a major impact on parasite motility. However, the motility of the double mutant parasites was indistinguishable from parasites expressing WT TgMLC1 in terms of motility initiation, mean displacement, mean speed and maximum speed (Fig. 6). Track length was slightly shorter in the C(8,11)S parasites, but still reached 88% of WT levels. Thus, parasites in which TgMLC1 has lost its ability to interact with TgGAP45 nevertheless show near normal motility.

**Figure 6.**
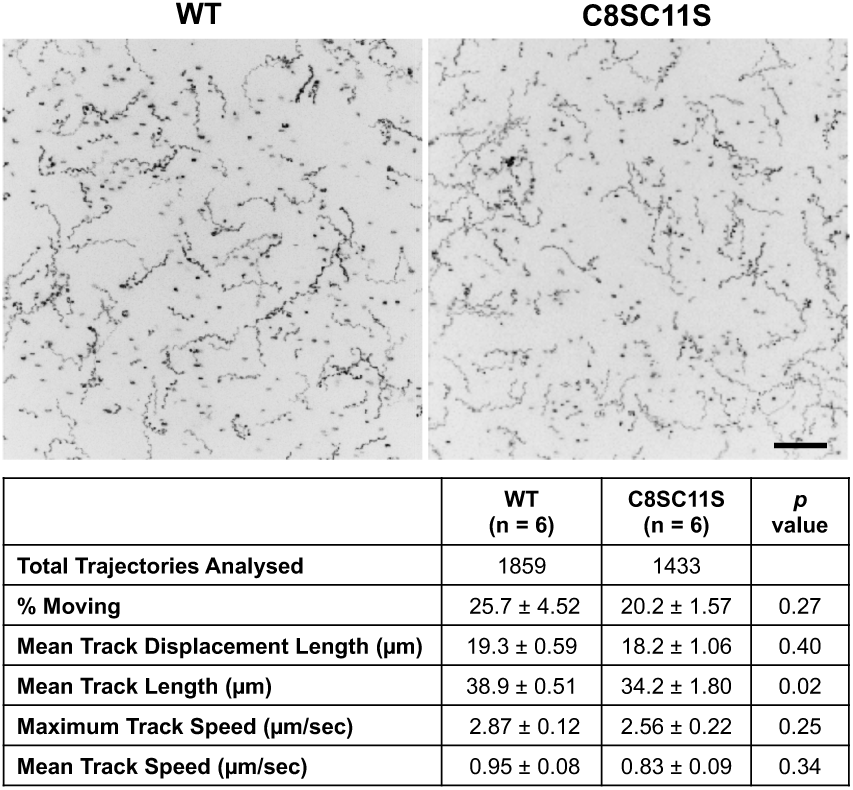
Mutations that block TgMLC1 palmitoylation and disrupt glideosome composition have little effect on parasite motility. The upper panels show maximum intensity projections of the Hoechst 33342-stained WT or C[8,11]S parasites that moved within a 3-dimensional model extracellular matrix (Matrigel) during 60 sec of image capture. Scale bar = 50 µm. The table below shows the motility parameters calculated from three independent motility assays (each consisting of three technical replicates); numbers for each parameter represent the means +/- SEM. The total number of trajectories analyzed for each parasite line is also shown. Differences between WT and C(8,11)S parasites for each motility parameter were assessed using an unpaired two-tailed t-test and the resulting *p* values are shown in the righthand column.

This result was unexpected since, according to the linear motor model of motility, disruption of the interaction between TgMLC1 and TgGAP45 should uncouple TgMyoA from the IMC, rendering it incapable of generating the force required for movement ((23); Fig. 1A). We therefore investigated whether the near normal motility observed in the mutants could be due to changes in the composition of the glideosome that could functionally compensate for the lack of TgMLC1-TgGAP45 interaction, such as the association of TgGAP45 with an alternative light chain/myosin motor, or interaction of either TgMLC1 or TgMyoA with alternative GAP proteins.

First, we asked whether TgGAP45 associates with any new proteins in the absence of its normal interaction with TgMLC1. Parasites expressing either WT or C(8,11)S TgMLC1 were metabolically labeled with ^35^S-methionine/cysteine and the labeled proteins that co-immunoprecipitated with TgGAP45 were resolved by SDS-PAGE, transferred to a PVDF membrane and visualized by phosphorimaging. The same membrane used for phosphorimaging was subsequently processed for western blotting with antibodies against TgMyoA, TgGAP45 and TgMLC1 to determine which of the ^35^S-labeled bands comigrate with which of the glideosome proteins. As expected, both TgMyoA and TgMLC1 co-immunoprecipitate with TgGAP45 from ^35^S-labeled WT parasites, whereas neither protein is recovered in TgGAP5 pulldowns from C(8,11)S parasites (Fig. 7A), confirming that the motor does not bind to TgGAP45 in the absence of TgMLC1 palmitoylation. No ^35^S-labeled bands were detected in the TgGAP45 pulldowns from C(8,11)S parasites that were not also present in the pulldowns from WT parasites (Fig. 7A). TgGAP45 therefore does not appear to interact to any significant/stoichiometric extent with alternate ^35^S-labeled myosins or myosin light chains when its interaction with TgMLC1 is disrupted by the C(8,11)S mutation.

**Figure 7.**
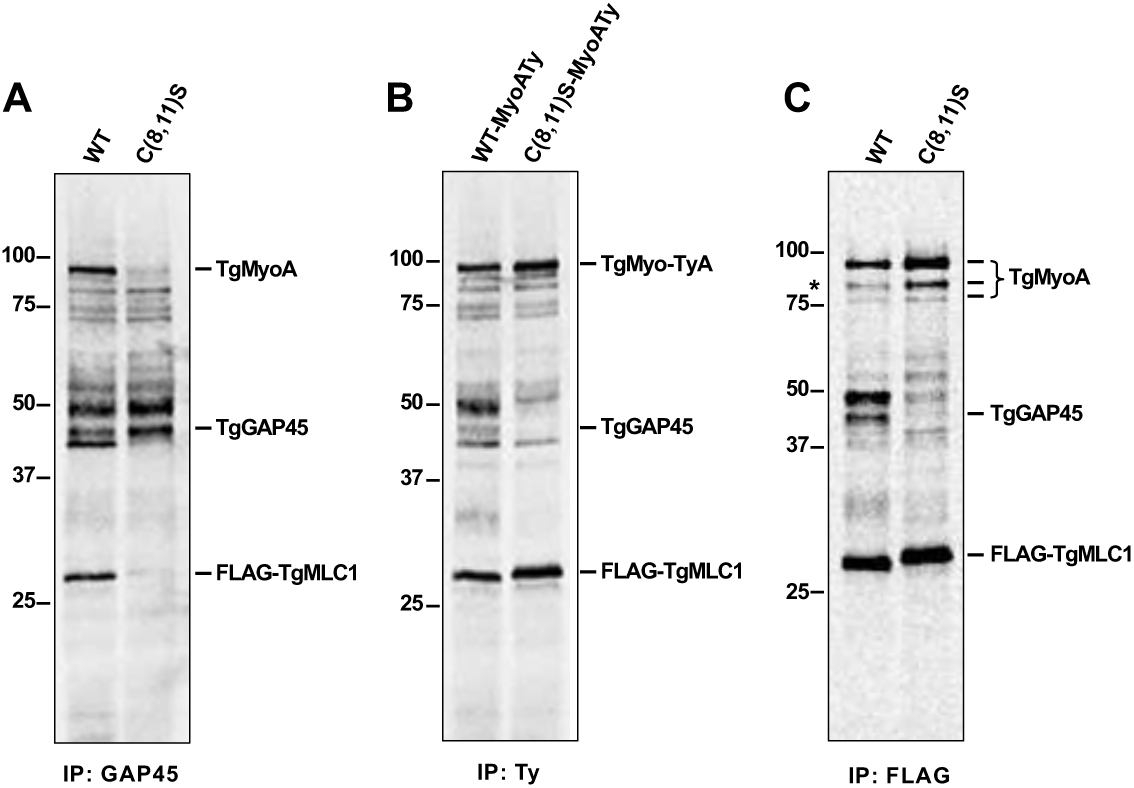
No changes in glideosome composition that could compensate for the loss of TgGAP45-TgMLC1 interaction are observed in C(8,11)S parasites. Parasites were labeled for 24 hr in medium containing (^35^S)-methionine/cysteine and gently extracted in Triton X-100. Soluble proteins were immunoprecipitated as described below, resolved by SDS-PAGE, transferred to a PVDF membrane and visualized by phosphorimaging (shown here). The same membrane was subsequently used for western blotting, probing with antibodies against TgMyoA, TgGAP45 and TgMLC1 (not shown) to determine which ^35^S-labeled bands corresponded to which glideosome proteins. Numbers on the left of each panel indicate molecular mass in kDa. **(A)** To test whether TgGAP45 interacts with any new proteins in C(8,11)S parasites, ^35^S-labeled proteins from WT and C(8,11)S parasites were immunoprecipitated with anti-TgGAP45, resolved by SDS/PAGE and visualized by phosphorimaging. No bands were detected in the pulldowns from C(8,11)S parasites that were not already present in the pulldowns from WT parasites. The band migrating immediately below TgGAP45 in the WT sample was recovered in some pulldowns but not others and may represent a breakdown product of TgGAPA45 (see Fig. S7). **(B)** To test whether TgMyoA interacts with any new proteins in the C(8,11)S parasites, ^35^S-labeled proteins immunoprecipitated using anti-Ty from WT-MyoATy and C(8,11)S-MyoATy parasites were compared. No bands were detected in the pulldowns from C(8,11)S parasites that were not already present in the pulldowns from WT parasites. **(C)** To test whether C(8,11)S TgMLC1 interacts with any new proteins compared to WT TgMLC1, ^35^S-labeled proteins immunoprecipitated using anti-FLAG from WT and C(8,11)S parasites were compared. Asterisk indicates an ∼80K band enriched in the FLAG pulldown of C(8,11)S compared to WT; this protein was shown to be a fragment of TgMyoA (see text). The relative amounts of TgMyoA and TgGAP45 pulled down in Panel C were quantified by western blotting as shown in Fig. S6.

We did a similar experiment to see if TgMyoA interacts with alternative GAP(s) (23) or other proteins in the C(8,11)S mutant, proteins that might serve to anchor TgMyoA into the IMC in the absence of a normal TgMLC1-TgGAP45 interaction. First, we Ty-tagged TgMyoA at the endogenous *TgMyoA* locus in both the WT and C(8,11)S backgrounds using CRISPR/Cas9 (Fig. S5). We then analyzed Ty pulldowns from ^35^S-labeled WT and C(8,11)S parasites. Ty-tagged TgMyoA and associated TgMLC1 were recovered in the Ty pulldowns from both parasite lines and, as expected, TgGAP45 was only recovered in pulldowns from WT parasites (Fig. 7B). Again, no ^35^S-labeled proteins were seen to associate with TgMyoA in the C(8,11)S parasites that were not also detected in WT parasites, so there is no evidence that TgMyoA interacts with new binding partners in the C(8,11)S double mutant (Fig. 7B).

Finally, we tested whether the mutant C(8,11)S TgMLC1 itself might interact with a different IMC-anchored protein(s) that could functionally compensate for its lack of interaction with TgGAP45. Anti-FLAG IPs from ^35^S-labeled WT and C(8,11)S parasites again confirmed the western blotting results presented in Figs. 5 and S3: compared to WT TgMLC1, C(8,11)S TgMLC1 shows both a greatly reduced interaction with TgGAP45 and an increased interaction with TgMyoA (Fig. 7C; quantitative western blot results from the same sample are shown in Fig. S6). However, in this experiment we also saw a marked increase in the amount of an ∼80kDa kDa ^35^S-labeled protein co-immunoprecipitating with C(8,11)S TgMLC1 compared to WT TgMLC1 (Fig 7C, asterisk). This was intriguing since TgGAP80, a TgGAP45-related protein that normally interacts with TgMyoC, is also reportedly capable of interacting with TgMLC1 (59). This raised the possibility that an increased interaction between C(8,11)S TgMLC1 and TgGAP80 might functionally compensate for the loss of interaction between TgMLC1 and TgGAP45 in the mutant parasites. We therefore repeated the FLAG-TgMLC1 pulldowns on a preparative scale and determined the identity of each of the major bands recovered by LC/MS/MS (Fig. S7 and Tables S1-S10). The ∼80kDa band proved to be a truncated form of TgMyoA rather than TgGAP80, and the increased levels of this TgMyoA fragment in the C(8,11)S pulldown paralleled the increased levels of full-length TgMyoA. Taken together, these data argue that the near normal motility seen in C(8,11)S parasites cannot be explained by the binding of either TgGAP45 or components of the motor to alternative proteins that could functionally compensate for the lack of interaction between TgMLC1 and TgGAP45.

## Discussion

The apicomplexan glideosome plays a critical role in parasite motility, invasion and virulence. Recent palmitome analyses have revealed that all known components of the *T. gondii* glideosome are palmitoylated, including TgMyoA, TgMLC1, TgELC1, TgGAP40, TgGAP45 and TgGAP50 (44, 54). Widespread glideosome palmitoylation has also been reported in *P. falciparum* (51, 60). Two of the *T. gondii* palmitoyl S-acyl transferases that are essential for parasite survival (TgDHHC2, TgDHHC14) localize to the IMC (61, 62) and are therefore well-situated to play a role in glideosome palmitoylation. The function of palmitoylation of one glideosome component, TgGAP45, was established in an elegant set of experiments by Frenal, Soldati-Favre and colleagues (23). Their study showed that C-terminal palmitoylation of TgGAP45 anchors its C-terminus in the IMC, while the other end of the protein is anchored in the plasma membrane via N-terminal palmitoylation and myristoylation. Acylation on the two ends of the protein therefore enables TgGAP45 to bridge the gap between the IMC and the plasma membrane; this determines the spacing between the two membranes and maintains the integrity of the parasite pellicle (23). The function of palmitoylation of other glideosome components is not known, and in most cases the specific residues palmitoylated on these other proteins have not been determined. We have focused here on the function of TgMLC1 palmitoylation.

In the linear motor model of motility, TgMLC1 plays two key roles in force generation: it stabilizes the TgMyoA lever arm (10, 13, 20), and it serves as a physical linker connecting the TgMyoA motor to TgGAP45 and the IMC (22, 23). We show here that TgMLC1 is dually palmitoylated, on C8 and C11 (Fig. 2). When we block palmitoylation by mutating these sites to either serine or alanine, TgMLC1 shows reduced association with the membrane fraction in phase-partitioning experiments (Fig. 4) and the mutant TgMLC1 no longer co-immunoprecipitates with TgGAP45 (Figs. 5, S3). Nevertheless, the non-palmitoylated protein continues to localize at the parasite periphery (Figs. 3, S1) and the motility of parasites expressing the mutant protein is in most aspects indistinguishable from wild-type parasites (Fig. 6).

How does palmitoylation of C8 and C11 affect the ability of TgMLC1 to bind TgGAP45? The detailed mechanism by which the N-terminus of TgMLC1 binds to the C-terminus of TgGAP45 (23) is unknown. Direct protein-protein interaction may be involved, as the mutation of two other sets of conserved N-terminal amino acids in TgMLC1 (D26E28 and P36GF38) also interfere with binding to TgGAP45 (23). The presence of the palmitates on C8 and C11 of TgMLC1 might help the interacting N-terminal residues of TgMLC1 transition from a disordered state (13, 20) into a binding-competent configuration. Alternatively, by inserting into the IMC membrane, the acyl chains could position the relevant N-terminal residues of TgMLC1 favorably for interaction with the C-terminus of TgGAP45, which is itself attached to the IMC membrane via palmitoylation. It is also possible that the acyl chains on TgMLC1 and TgGAP45 themselves interact within the plane of the membrane (20). In any of these cases, blocking TgMLC1 palmitoylation would be expected to inhibit TgMLC1-TgGAP45 interaction, as observed. It is unlikely that palmitoylation is necessary for proper folding and stability of TgMLC1, as appears to be the case with the *P. falciparum* MLC1 homolog, MTIP (51), since (a) we see no evidence for increased degradation of the C(8,11)S mutant compared to WT TgMLC1 (Fig. S4) and (b) the palmitoylation-deficient mutants continue to bind to TgMyoA (*e*.*g*., Fig 5).

Our data argue against the model that binding to TgGAP45 is what localizes TgMLC1 (and, therefore, TgMyoA) to the parasite periphery and anchors the motor in the IMC membrane. We showed that palmitoylation-deficient TgMLC1 no longer interacts with TgGAP45, yet both TgMLC1 (Figs. 3, S1) and TgMyoA (Fig. S5) continue to localize to the periphery in these mutant parasites. Previous studies with parasites expressing mutant TgGAP45 also argue against a model in which the localization of TgMLC1 is determined by TgGAP45: TgGAP45 lacking its C-terminal palmitoylation sites becomes dissociated from the IMC, yet TgMLC1 nevertheless remained IMC associated (23). It is not clear how palmitoylated TgMLC1 located within the space between the IMC and plasma membrane would be targeted to the IMC rather than the plasma membrane, if not through TgGAP45. The simplest hypothesis is that it interacts directly with one of the other resident proteins of the IMC, such as the transmembrane proteins TgGAP40 or TgGAP50. Alternatively, insertion of TgMLC1 into the IMC could be a direct consequence of binding to (63) and/or palmitoylation by IMC-localized palmitoyl S-acyl transferases (61, 62).

The most unexpected result of this study was that the motility of parasites expressing C(8,11)S TgMLC1 was in most aspects indistinguishable from wild-type parasites (Fig. 6). This observation suggests that the coupling of the motor complex to the IMC via TgGAP45 is unnecessary for force generation by the parasite, and directly contradicts a fundamental tenet of the linear motor model of motility. The near normal motility observed in the mutants does not appear to be due to compensatory changes in glideosome composition in response to the mutations (*i*.*e*., TgGAP45 does not associate with alternative myosin light chain(s) or myosin motor(s) in the C[8,11]S mutant, nor does the mutant TgMLC1 or TgMyoA interact with alternative GAP proteins; Fig. 7). It remains formally possible that non-palmitoylatable TgMLC1 binds to TgGAP45 in the parasite, with a reduced affinity that is sufficient to maintain its association with the IMC and support motility but insufficient to survive detergent extraction and immunoprecipitation. While it is difficult to experimentally exclude this possibility, most previous studies of glideosome composition and function have utilized a similar nonionic detergent extraction/immunoprecipitation approach, which constitutes the operational definition of the glideosome (17, 18, 23). Furthermore, our phase partitioning experiments independently suggest that the mutant TgMLC1 has indeed lost its association with the IMC membrane. Finally, recent data demonstrate that parasites lacking TgMLC1 altogether have a defect in motility initiation, but those parasites that continue to move do so at normal speeds (25).

At a minimum, our data demonstrate that TgMLC1 palmitoylation affects its binding to TgGAP45 but this palmitoylation plays little to no role in parasite motility, as assayed by the most sensitive and quantitative assays currently available. The data also show clearly that the inhibitory effects of the palmitoylation inhibitor 2-bromopalmitate on parasite motility are not due to changes in palmitoylation of TgMLC1, as previously hypothesized, although changes in the palmitoylation of other glideosome components could be involved (64). While the data presented here do not, by themselves, disprove the linear motor model of motility, they add to a growing list of evidence (24-29, 33) suggesting that the mechanisms underlying apicomplexan parasite motility are more complicated than what is currently encapsulated by the linear motor model. How the different motility-associated proteins of the parasite interact and work together to generate the forces necessary to drive parasite movement and whether more than one underlying mechanism exists remain important open questions for future study.

## Materials and Methods

### Parasite culture

*T. gondii* tachyzoites were maintained by serial passage in confluent monolayers of human foreskin fibroblasts (HFFs) (ATCC CRL-1634) grown in Dulbecco’s Modified Eagle’s Medium (DMEM), supplemented with 10% (v/v) heat-inactivated fetal bovine serum (FBS) and 10 mM HEPES, pH 7.0, as previously described (65). The medium was changed to DMEM supplemented with 1% (v/v) heat-inactivated FBS and 10 mM HEPES pH 7.0 prior to infection of confluent HFFs with parasites.

### Generation of TgMLC1 knock-in mutants by allelic replacement

Mutations were introduced into a previously described *TgMLC1* allelic replacement plasmid (55) using the Quick Change site-directed mutagenesis kit (Agilent Technologies). *E*.*coli* were transformed with the mutagenized plasmids and colonies screened by colony PCR and restriction digestion. The entire open reading frame was sequenced to confirm the presence of only the desired mutation(s). The allelic replacement plasmid was linearized with BglII and PciI and used to transfect RHΔ*ku80*Δ*HXGPRT* parasites. Successful integration of the mutated gene at the endogenous locus yields phleomycin-resistant, FLAG-positive parasites. Parasites were therefore subjected to two rounds of phleomycin selection, cloned by limiting dilution and characterized by anti-FLAG immunofluorescence and immunoblotting, as well as diagnostic PCR to confirm correct integration on the chromosome (see Fig. S1). Finally, the presence of the desired mutations in individual clones was confirmed by sequencing of genomic DNA.

### Labeling with 17-ODYA

ODYA labeling was performed as described previously (54).

### Epitope tagging of the *TgMyoA* locus using CRISPR/Cas9

To insert the coding sequence for a Ty epitope tag at the C-terminus of *TgMyoA*, we constructed plasmid pU6-MyoA, which contains the *TgMyoA* targeting chiRNA under the U6 promoter and *Cas9* under the *TUB1* promoter (66). First, we synthesized a dsDNA oligo encoding the protospacer sequence used to direct Cas9 to the C-terminal region of *TgMyoA*. To fuse the *TgMyoA* protospacer to the chiRNA of the pU6-universal plasmid, forward and reverse oligos corresponding to the *TgMyoA* 3’ region were annealed by combining them (20μl each, 200μM stocks) in duplex buffer (100mM potassium acetate; 30mM HEPES pH 7.5), heating them to 100°C for 2 minutes, then slowly cooling them to 25°C and letting them stand overnight to generate double-stranded product. The duplexed oligos were dialyzed against deionized water, phosphorylated using T4 polynucleotide kinase, and heat inactivated. Meanwhile, the pU6-universal plasmid (5μg) was linearized by digesting with BsaI, dephosphorylated with Antarctic phosphatase, heat inactivated and PCR purified. The phosphorylated, duplexed oligos were then ligated into the pU6-universal plasmid to generate pU6-MyoAPS. Competent *E. coli* DH5-*α* were transformed with the pU6-MyoAPS ligation mixture, and individual colonies with the desired plasmid identified first by colony PCR, then by diagnostic PCR and sequencing. pU6-MyoAPS was transfected along with duplexed, dialyzed homologous recombination (HR) oligos into WT and C(8,11)S parasites. Ty-positive parasites were identified by immunofluorescence, cloned by limiting dilution and confirmed as expressing TgMyoA-Ty from the endogenous *TgMyoA* locus by immunofluorescence, diagnostic PCR (see Fig. S5), and sequencing of genomic DNA.

### Immunofluorescence

HFF cells were infected for 15h and fixed with 4% (v/v) paraformaldehyde in PBS (15 min, 25°C). Fixed cells were washed and permeabilized with PBS containing 0.25% (v/v) Triton-X-100 for 20 minutes, washed 3 times with PBS and blocked (30 min, 25°C) in Block buffer (PBS containing 1% [w/v] BSA). The cells were then incubated for 1hr with primary antibodies diluted as follows with Block buffer: mouse monoclonal anti-FLAG (Sigma Aldrich) at 1:500; rabbit anti-GAP45 polyclonal serum (a generous gift from Dr. Con Beckers (17)) at 1:1000; rabbit anti-MyoA polyclonal serum (67) at 1:20; and rabbit anti-ELC1 polyclonal serum at 1:500. Samples were washed 3 times and incubated a further 30 min in goat-anti-rabbit IgG conjugated to Alexa fluor 546 (Invitrogen) or goat-anti-mouse IgG conjugated to Alexa fluor 488 (Invitrogen), each diluted 1:500 in Block buffer. After four final washes in PBS, fluorescence was visualized by epifluorescence microscopy.

### Immunoprecipitation

For anti-FLAG immunoprecipitations, 2×10^7^ freshly egressed parasites were extracted for 45 minutes on ice in 3 ml of FLAG lysis buffer (10 mM imidazole pH 7.4, 300 mM NaCl, 1 mM EGTA, 5 mM MgCl_2_, 1 % (w/v) TX-100, 2 mM DTT, 2mM ATP) containing 1:100 (v/v) protease inhibitors (Sigma #P8340). The extract was divided into two equal portions and insoluble material was pelleted at 10,000*xg* (30 min, 4°C). Each ∼1.5 ml of supernatant was used to resuspend 20 ul of packed anti-FLAG M2 affinity resin (Sigma), then rocked gently overnight at 4°C. After three washes with the FLAG wash buffer (FLAG lysis buffer containing 1:500 (v/v) protease inhibitors), bound proteins from the pooled washed resin were eluted with 100μg of FLAG peptide (Sigma) in FLAG wash buffer. Eluates were resolved by SDS-PAGE and proteins visualized by immunoblotting. Primary and secondary (IRDye 680-conjugated anti-rabbit IgG and IRDye 800-conjugated anti-mouse IgG) antibodies were diluted for use in Odyssey blocking buffer (LI-COR). The blots were scanned using an Odyssey CLx infrared imager (LI-COR). Images were processed using Image Studio software (LI-COR). Signal intensities of bands being compared were normalized as described in the figure legends.

### ^35^S-Metabolic labeling

For ^35^S metabolic labeling, confluent HFF cells in a T75 flask were infected with 1×10^7^ tachyzoites and incubated for 16-20 hours. The infected cells were then incubated in methionine/cysteine-free DMEM (GIBCO) containing 1% (v/v) FBS for 1 hr, followed by 24 hr in DMEM containing 500µCi ^35^S-Easytag mix (Perkin Elmer). Infected cells were detached from the flask using a cell scraper, washed twice with ice-cold PBS, and lysed in FLAG lysis buffer for anti-FLAG immunoprecipitation, or TX-100 lysis buffer (1% v/v TX-100, 50 mM Tris HCl pH 8.0, 150 mM NaCl, 2 mM EDTA and 1:200 (v/v) protease inhibitors) for anti-Ty and anti-GAP45 immunoprecipitations. Immunoprecipitation was performed as described above except that, after incubation with primary antibody, protein A-Sepharose (Invitrogen) was added and incubated for 1hr with gentle agitation at 4°C to collect the immune complexes. After three washes with either FLAG wash buffer or TX-100 IP wash buffer (1% v/v TX-100, 50 mM Tris, pH 8.0, 150 mM NaCl, 5 mM EDTA and 1:500 protease inhibitors), bound proteins were eluted in SDS-PAGE sample buffer by boiling at 100°C for 5 minutes. Samples were then resolved by SDS-PAGE, and transferred to PVDF membranes for phosphorimaging and immunoblotting.

### Phase separation of parasite proteins in Triton X-114

The phase separation was performed as previously described (56-58). Briefly, 4×10^8^ tachyzoites were extracted in 1 ml extraction buffer (10 mM Tris-HCl, pH 7.4, 150 mM NaCl, 0.5% (v/v) precondensed TX-114 (Pierce) and 1:100 [v/v] dilution of protease inhibitors) for 90 minutes on ice. Insoluble material was removed by centrifugation (twice at 13,000x*g* for 5 min at 4°C). The cleared extract was overlaid onto a 750 μl prechilled sucrose cushion (6% [w/v] sucrose, 10 mM Tris-HCl, pH 7.4, 150 mM NaCl and 0.06% [v/v] precondensed Triton X-114), incubated for 5 minutes at 37°C and centrifuged to separate the detergent and aqueous phases by centrifugation (37°C, 3000x*g*, 5 min). Partitioning was repeated once on each phase and the collected samples were resolved by SDS-PAGE and analyzed by sequential incubations of a single western blot with anti-TgMyoA, -TgGAP45, -TgMLC1 and -TgELC1, followed by the appropriate secondary antibodies.

### Motility assays

3D motility assays in Matrigel were performed as previously described (55, 68).

### Protein identification by mass spectrometry analysis

Dried tryptic peptides recovered from excised gel bands (69) were dissolved in 10 µl 0.1% formic acid and 2.5% acetonitrile, and 2 ul were analyzed on the Thermo Q-Exactive mass spectrometer coupled to an EASY-nLC system (Thermo Fisher). Peptides were separated on a fused silica capillary (12 cm x 100 um I.D) packed with Halo C18 (2.7 um particle size, 90 nm pore size, Michrom Bioresources) at a flow rate of 300 nl/min. Peptides were introduced into the mass spectrometer via a nanospray ionization source at a spray voltage of 2.2 kV. Mass spectrometry data were acquired in a data-dependent top-10 mode, and the lock mass function was activated (m/z, 371.1012). Full scans were acquired from m/z 350 to 1,600 at 70,000 resolution (automatic gain control [AGC] target, 1e6; maximum ion time [max IT], 100 ms; profile mode). Resolution for dd-MS^2^ spectra was set to 17,500 (AGC target: 1e5) with a maximum ion injection time of 50 ms. The normalized collision energy was 27 eV. A gradient of 0 to 40% acetonitrile (0.1% FA) over 55 min was applied. The spectra were searched against the *T. gondii* protein database (http://www.toxodb.org/toxo/) by Proteome Discoverer (PD) 2.0. The search parameters permitted a 10 ppm precursor MS tolerance and a 0.02 Da MS/MS tolerance. Carboxymethylation of cysteines was set up as fixed modification and Oxidation of methionine (M) was allowed as variable modification. Up to three missed tryptic cleavages of peptides were considered with the false-discovery rate set to 1% at the peptide level.

## Figure and Table Legends

**Table 1:** Strains generated and used in this study, with their relevant genotypes.

| Strain designation | Relevant genotype |
| --- | --- |
| WT | *RHΔku80Δmlc1::Flag-MLC1* |
| C8S | *RHΔku80Δmlc1::Flag-MLC1<sup>C8S</sup>* |
| C11S | *RHΔku80Δmlc1::Flag-MLC1<sup>C11S</sup>* |
| C(8,11)S | *RHΔku80Δmlc1::Flag-MLC1<sup>C8SC11S</sup>* |
| C(8,11)A | *RHΔku80Δmlc1::Flag-MLC1<sup>C8AC11A</sup>* |
| WT-MyoATy | *RHΔku80Δmlc1::Flag-MLC1ΔmyoA::MYOA-Ty* |
| C(8,11)S-MyoATy | *RHΔku80Δmlc1::Flag-MLC1<sup>C8SC11S</sup>ΔmyoA::MYOA-Ty* |

## Supplementary figure legends and tables

**Figure S1.**
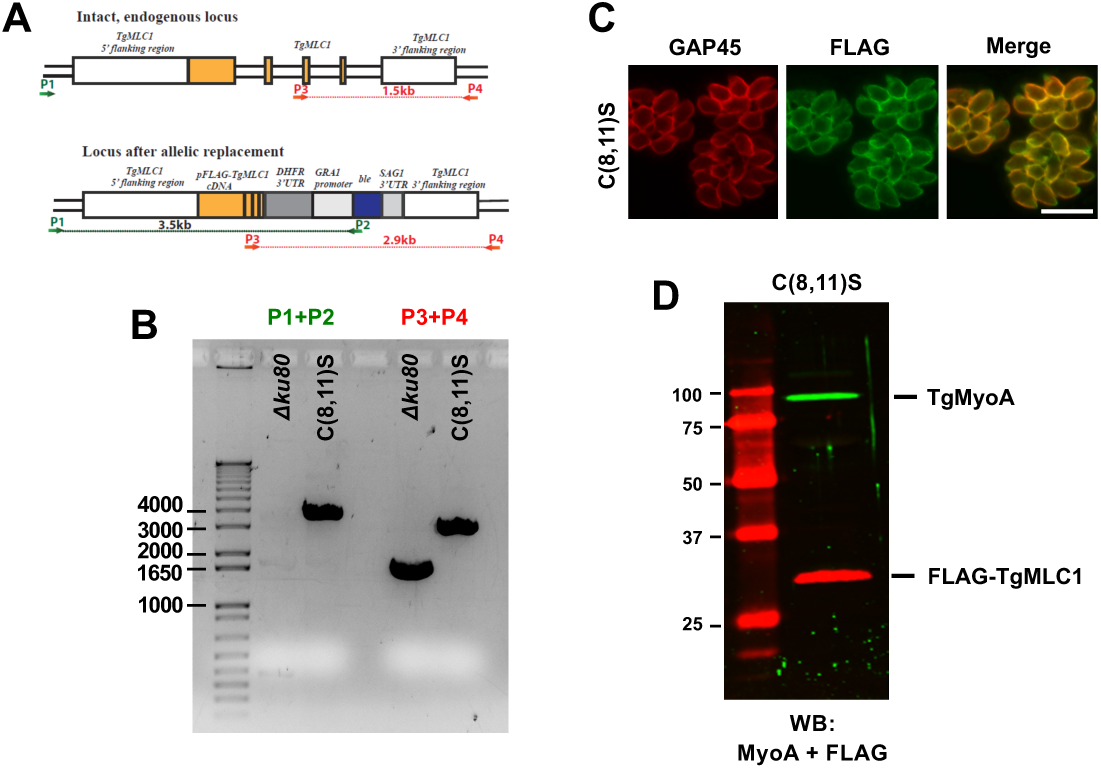
Generation and characterization of TgMLC1 knock-in parasites. **(A)** Allelic replacement strategy. The *TgMLC1* allele was targeted in *RHΔku80* parasites using 5’ and 3’ *TgMLC1* homology regions flanking a construct that consisted of cDNA encoding full-length TgMLC1 (wildtype or mutant) with an N-terminal FLAG tag, the *3’* untranslated region (UTR) from *DHFR*, and a selection cassette consisting of the *GRA1* promoter, the phleomycin resistance gene (*ble*) and the 3’ UTR from *SAG1*. The binding location of PCR primers P1-P4 (which correspond to Primers 25-28, respectively, in Supplementary Table 11) and the corresponding expected amplicon sizes are indicated in red. This same strategy was used to generate all TgMLC1 knock-in lines. Panels B-D illustrate how insertion at the genomic locus and correct localization and expression of the mutant protein was confirmed in each knock-in line; results for the C(8,11)S parasite line are shown here. **(B)** PCR results using primer pairs P1+P2 and P3+P4 on parental (*Δku80)* and C(8,11)S parasites after allelic replacement. DNA ladder is shown in the leftmost lane, numbers indicate base pairs. **(C)** Immunofluorescence staining of C(8,11)S parasites with anti-GAP45 (red) and anti-FLAG (green) confirms expression and proper (peripheral) localization of the FLAG-tagged TgMLC1. Scale bar = 10 μm. **(D)** Western blot of C(8,11)S parasites stained with anti TgMyoA (green) and anti-FLAG (red) confirms expression and proper size of the FLAG-tagged protein. Numbers on the left indicate molecular mass in kDa. Successful allelic replacement by constructs expressing WT, C8S, C11S and C8A11A TgMLC1 were similarly confirmed by PCR, immunofluorescence and western blot.

**Figure S2.**
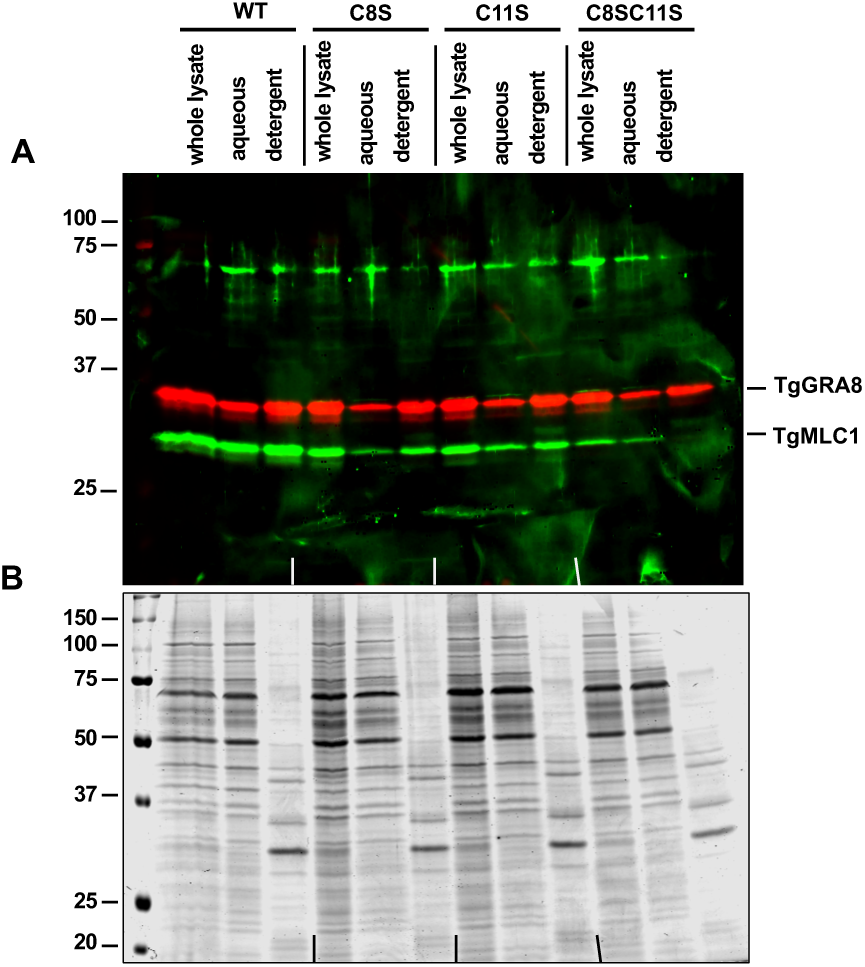
Triton X-114 phase partitioning of parasites expressing WT *vs*. palmitoylation-deficient TgMLC1. WT, C8S, C11S and C8SC11S parasites were extracted at 4°C in Triton X-114 and subjected to phase partitioning at 20°C. Proteins present in the lysate before partitioning (Whole lysate) and in the aqueous and detergent phases after partitioning were resolved by SDS-PAGE and visualized either by **(A)** Western blotting with anti-TgMLC1 (green) and anti-TgGRA8 (red), or **(B)** Colloidal Coomassie staining. Numbers on the left of each panel indicate molecular mass in kDa.

**Figure S3.**
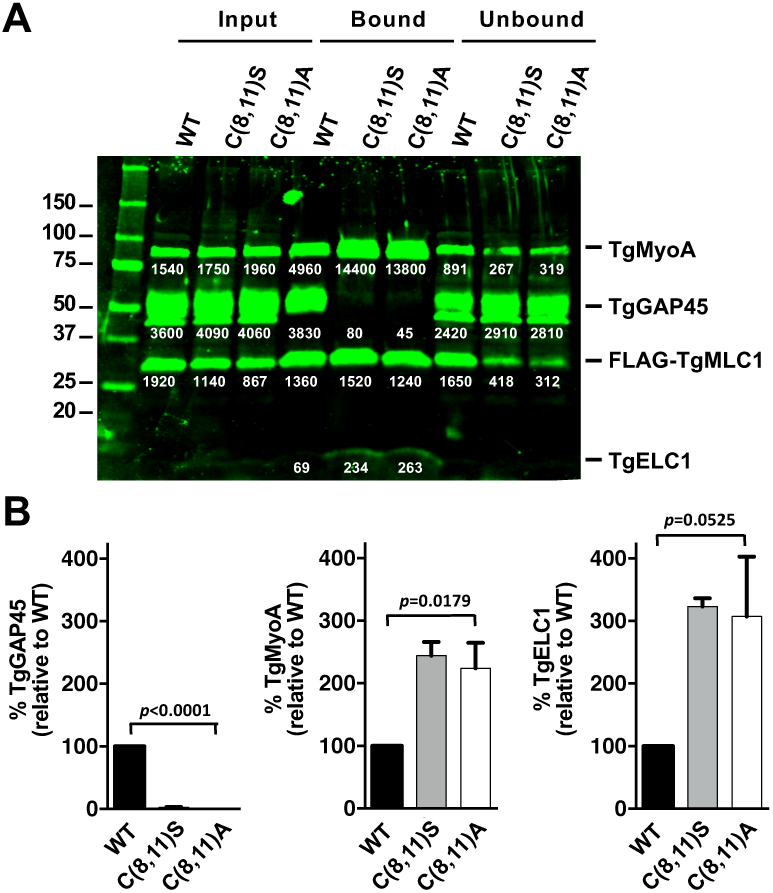
Similar changes in the composition of the glideosome are observed in C(8,11)S and C(8,11)A parasites. **(A)** Parasites expressing wild-type or mutant FLAG-tagged TgMLC1 were gently extracted in Triton X-100, and the soluble proteins (Input) used for anti-FLAG immunoprecipitations. Proteins that bound to the anti-FLAG affinity resin and those that did not (Bound, Unbound) were recovered and analyzed by SDS-PAGE and sequential western blotting with anti-TgMyoA, -TgGAP45, -TgMLC1 and -TgELC1. The signal intensity of each immunoreactive band was quantified (white number below the band; relative fluorescence units). Numbers on the left indicate molecular mass in kDa. **(B)** The signal intensities of TgGAP45, TgMyoA and TgELC1 pulled down from each of the parasite lines is shown relative the corresponding band from WT parasites, after normalizing to the amount of TgMLC1 recovered in each sample. Shown are the means and SEM from three independent experiments; data were analyzed using one-way ANOVA.

**Figure S4.**
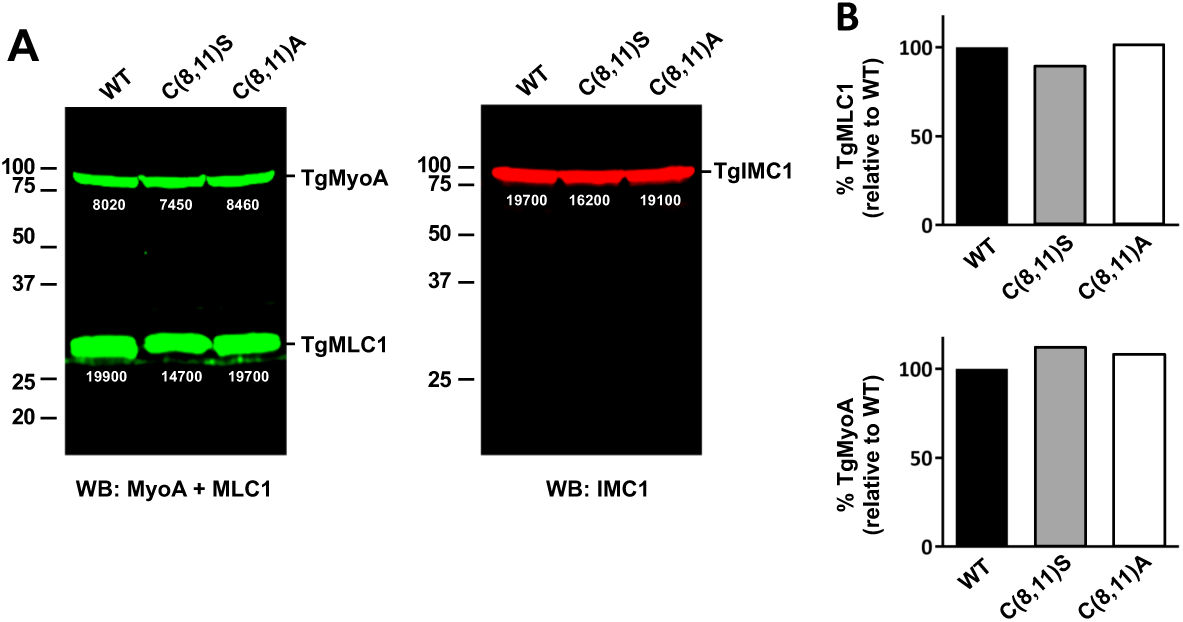
WT, C(8,11)S and C(8,11)A parasites express equivalent amounts of both FLAG-tagged TgMLC1 and TgMyoA. Parasite proteins were extracted in boiling SDS-PAGE sample buffer, resolved by SDS-PAGE and visualized by western blotting with anti-TgMyoA and anti-TgMLC1 (left panel) or anti-TgIMC1 (right panel) as a loading control. The signal intensity of each immunoreactive band was quantified (white number below the band; relative fluorescence units). Numbers on the left indicate molecular mass in kDa. **(B)** The signal intensities of TgMyoA and TgMLC1 in each of the parasite lines is shown relative the corresponding band from wildtype parasites, after normalizing to the amount of TgIMC1 in each sample.

**Figure S5.**
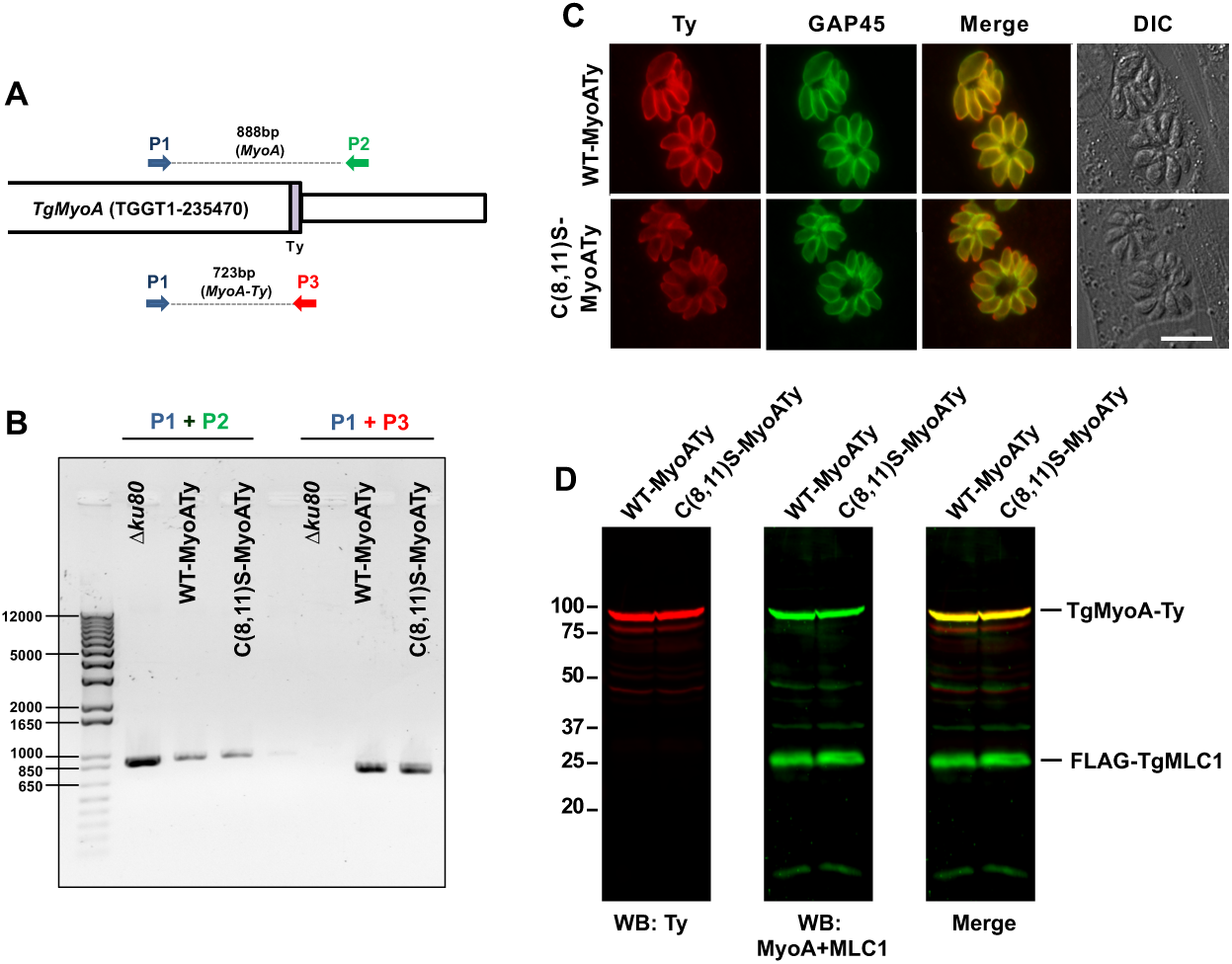
Generation and characterization of parasites expressing C-terminally Ty-tagged TgMyoA. **(A)** Schematic representation of the *TgMyoA* locus after the insertion of Ty tag at the 3’ end using CRISPR-Cas9. The binding location of PCR primers P1-P3 and the expected amplicon sizes are indicated. Primers P1, P2 and P3 correspond to primers 20, 22 and 21, respectively, in Supplementary Table 11. **(B)** PCR results using primer pairs P1+P2 and P1+P3 on parental parasites (*Δku80)* and the WT and C(8,11)S lines after Ty tagging the *TgMyoA* locus. DNA ladder is shown in the leftmost lane, numbers indicate base pairs. **(C)** Immunofluorescence staining of WT-MyoATy and C(8,11)S-MyoATy parasites with anti-Ty (red) and anti-GAP45 (green) confirms expression and proper localization of Ty-tagged TgMyoA in the two lines. Scale bar = 10 μm. **(D)** Western blots of WT-MyoATy and C(8,11)S-MyoATy parasites probed with anti-Ty (red) followed by anti-TgMyoA and TgMLC1 (green) confirms similar expression levels of TgMyoA in the two lines. Numbers on the left indicate molecular mass in kDa.

**Figure S6:**
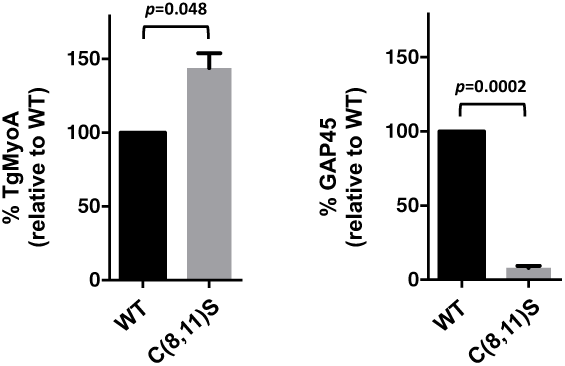
The western blot signal intensities of **(A)** TgMyoA and **(B)** TgGAP45 pulled down from WT or C(8,11)S parasites in Fig. 7C, after normalizing for the amount of TgMLC1 recovered in each sample. Shown are the means and SEM from two independent experiments; differences were assessed using an unpaired two-tailed t-test.

**Figure S7.**
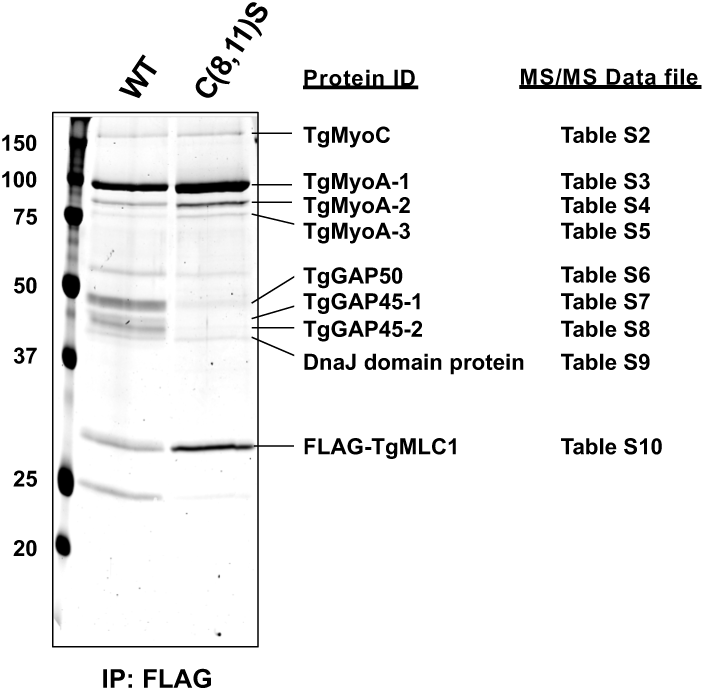
Preparative SDS-PAGE of anti-FLAG immunoprecipitated proteins from WT and C(8,11)S parasites. Proteins immunoprecipitated using anti-FLAG from WT and C(8,11)S parasites were resolved by SDS-PAGE and stained with colloidal Coomassie Blue. The major stained bands were excised, digested with trypsin and identified by LC/MS-MS as described in Materials and Methods. Numbers on the left indicate molecular mass in kDa. Labels on the right indicate the major protein identified in the band and the LC/MS-MS data file on which this identification was based (see Table S1 for a summary of the LC/MS-MS results and Tables S2-S10 for the raw data for each band).

**Supplementary Tables 1-10:**
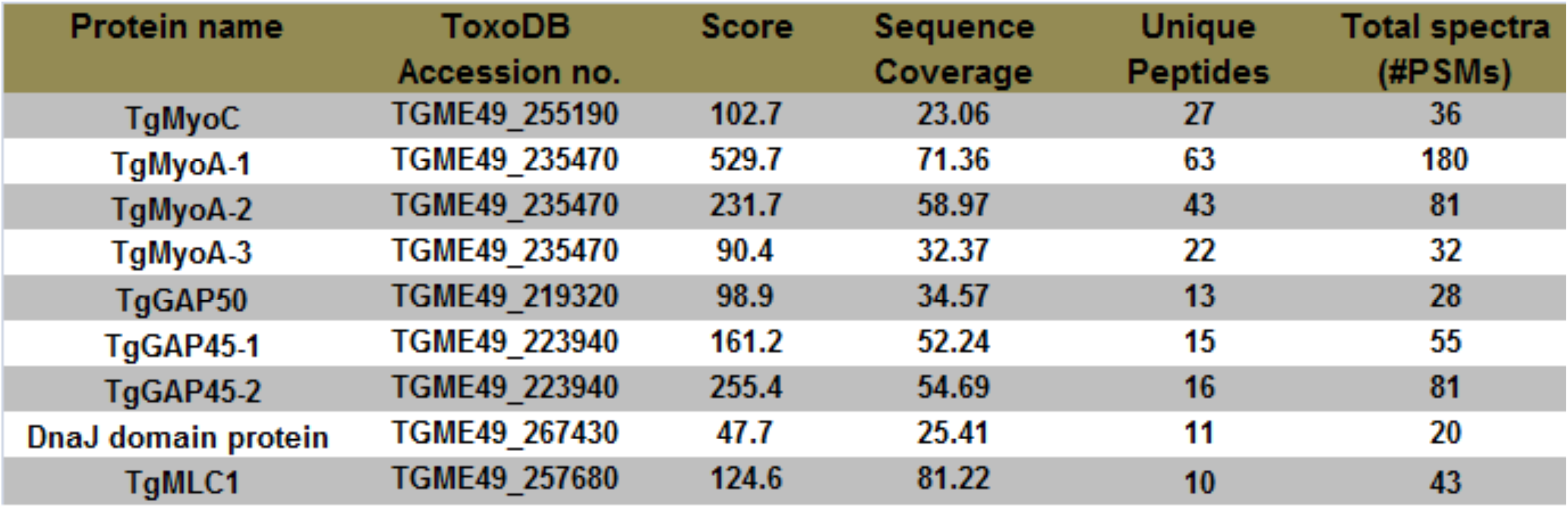
Identification of protein bands recovered in the anti-FLAG immunoprecipitates from WT and C(8,11)S parasites. Protein identification data for the nine excised bands shown in Fig. S6 are summarized in Table S1, and the raw LC/MS-MS results for each individual band are provided in Tables S2-S10.

**Supplementary Tables 1-10:**
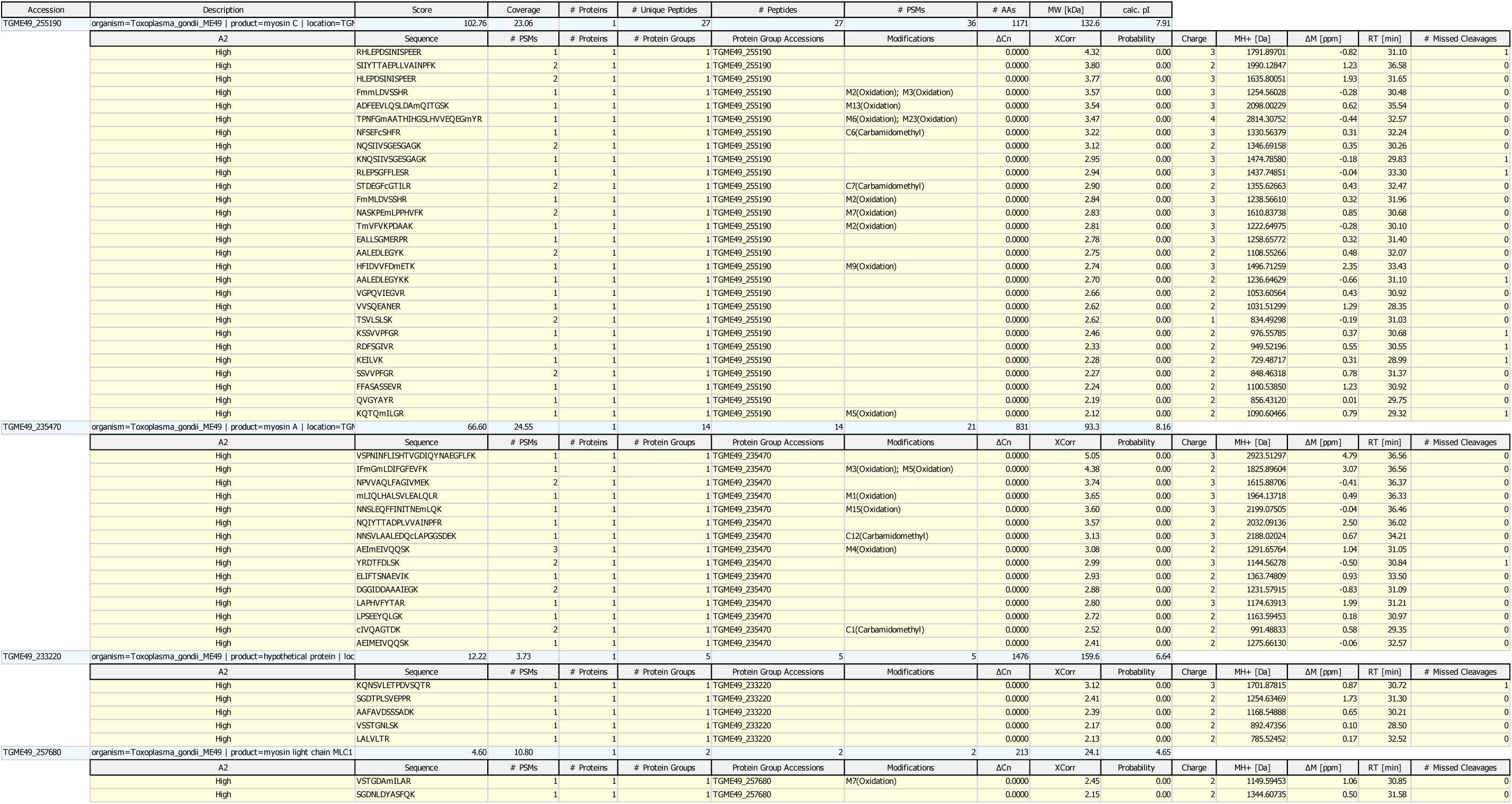
Identification of protein bands recovered in the anti-FLAG immunoprecipitates from WT and C(8,11)S parasites. Protein identification data for the nine excised bands shown in Fig. S6 are summarized in Table S1, and the raw LC/MS-MS results for each individual band are provided in Tables S2-S10.

**Supplementary Tables 1-10:**
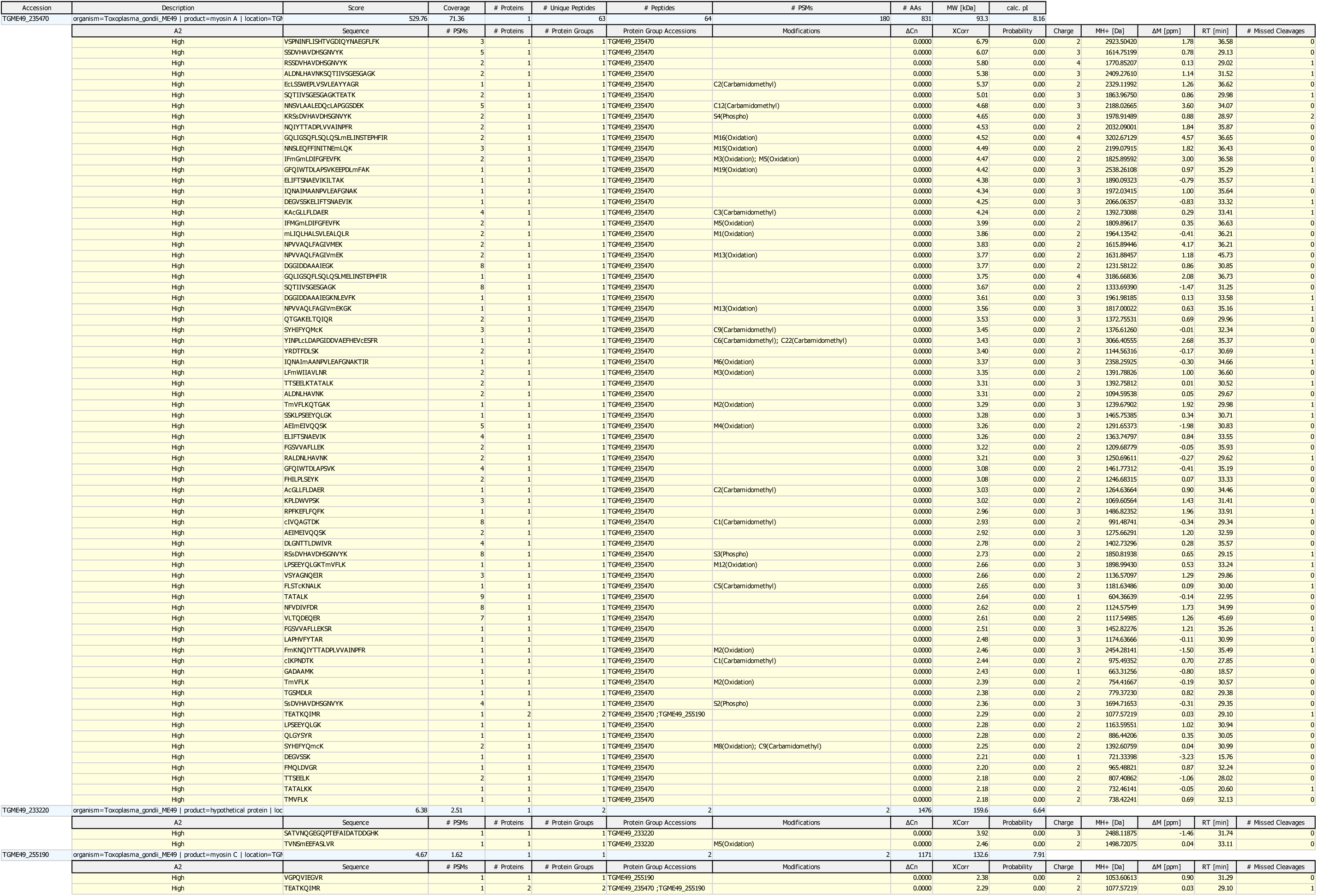
Identification of protein bands recovered in the anti-FLAG immunoprecipitates from WT and C(8,11)S parasites. Protein identification data for the nine excised bands shown in Fig. S6 are summarized in Table S1, and the raw LC/MS-MS results for each individual band are provided in Tables S2-S10.

**Supplementary Tables 1-10:**
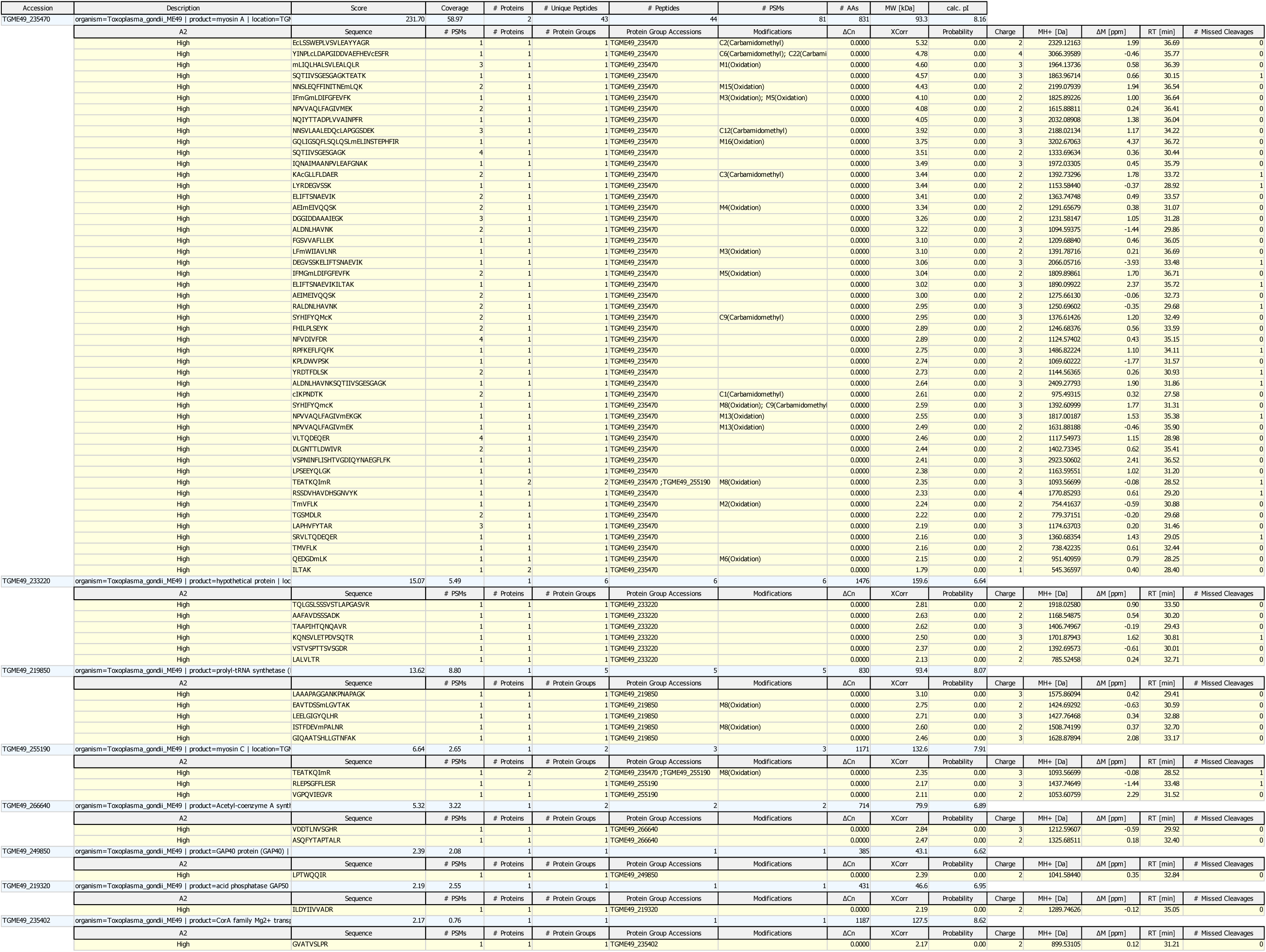
Identification of protein bands recovered in the anti-FLAG immunoprecipitates from WT and C(8,11)S parasites. Protein identification data for the nine excised bands shown in Fig. S6 are summarized in Table S1, and the raw LC/MS-MS results for each individual band are provided in Tables S2-S10.

**Supplementary Tables 1-10:**
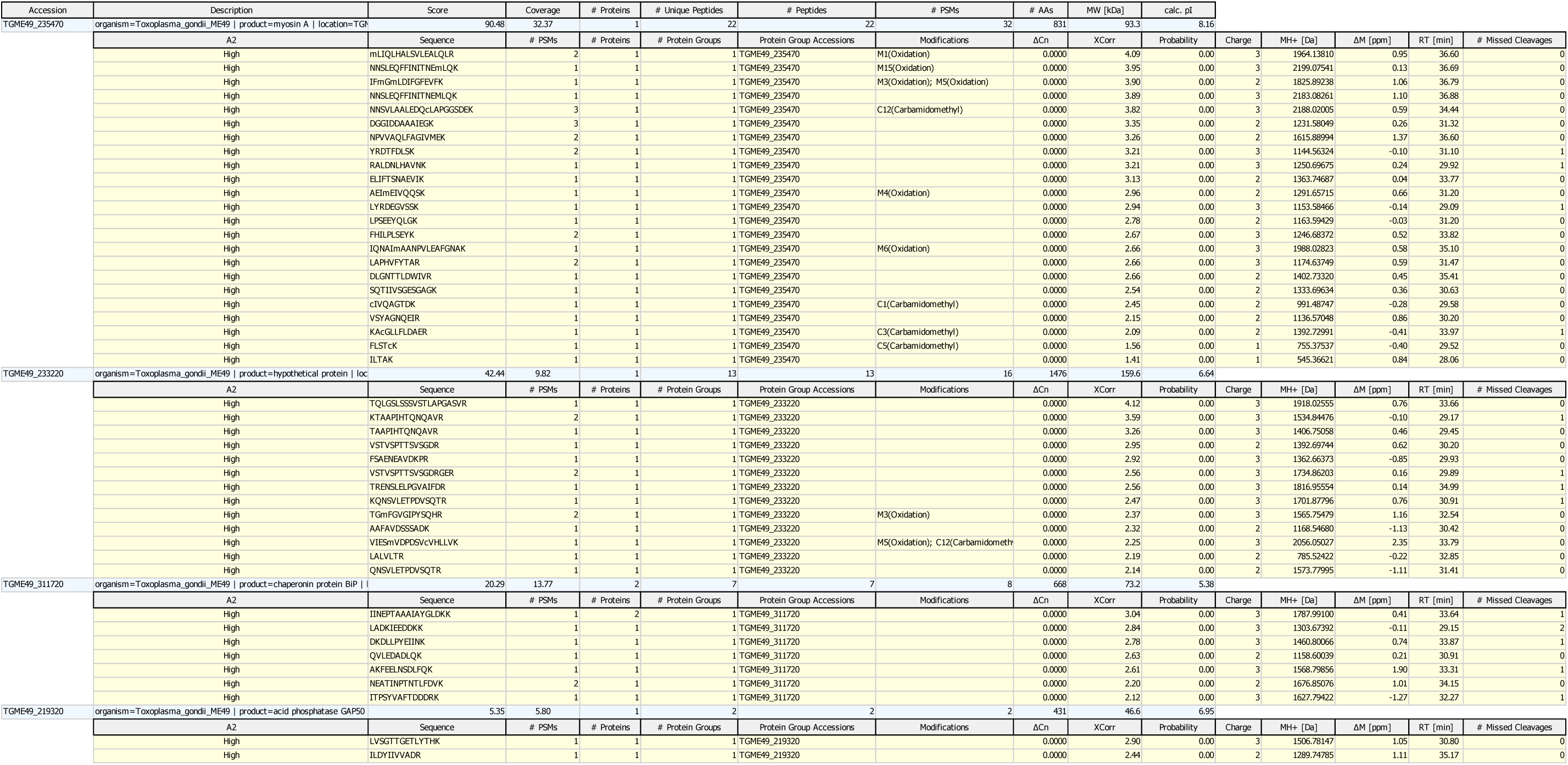
Identification of protein bands recovered in the anti-FLAG immunoprecipitates from WT and C(8,11)S parasites. Protein identification data for the nine excised bands shown in Fig. S6 are summarized in Table S1, and the raw LC/MS-MS results for each individual band are provided in Tables S2-S10.

**Supplementary Tables 1-10:**
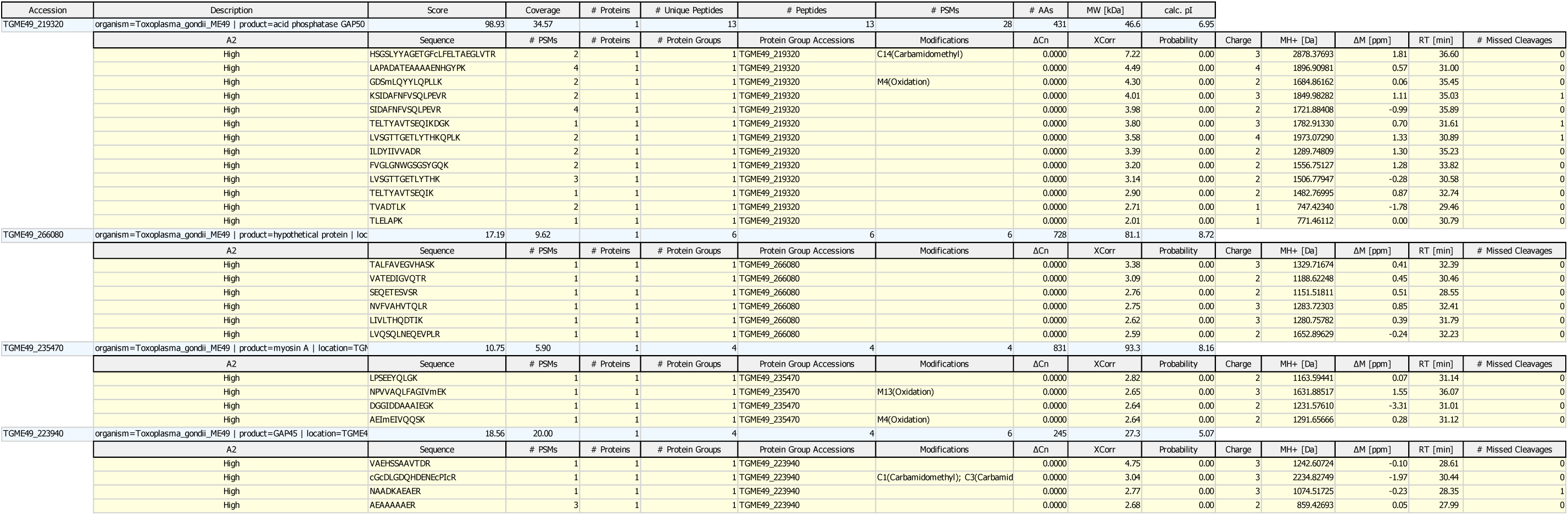
Identification of protein bands recovered in the anti-FLAG immunoprecipitates from WT and C(8,11)S parasites. Protein identification data for the nine excised bands shown in Fig. S6 are summarized in Table S1, and the raw LC/MS-MS results for each individual band are provided in Tables S2-S10.

**Supplementary Tables 1-10:**
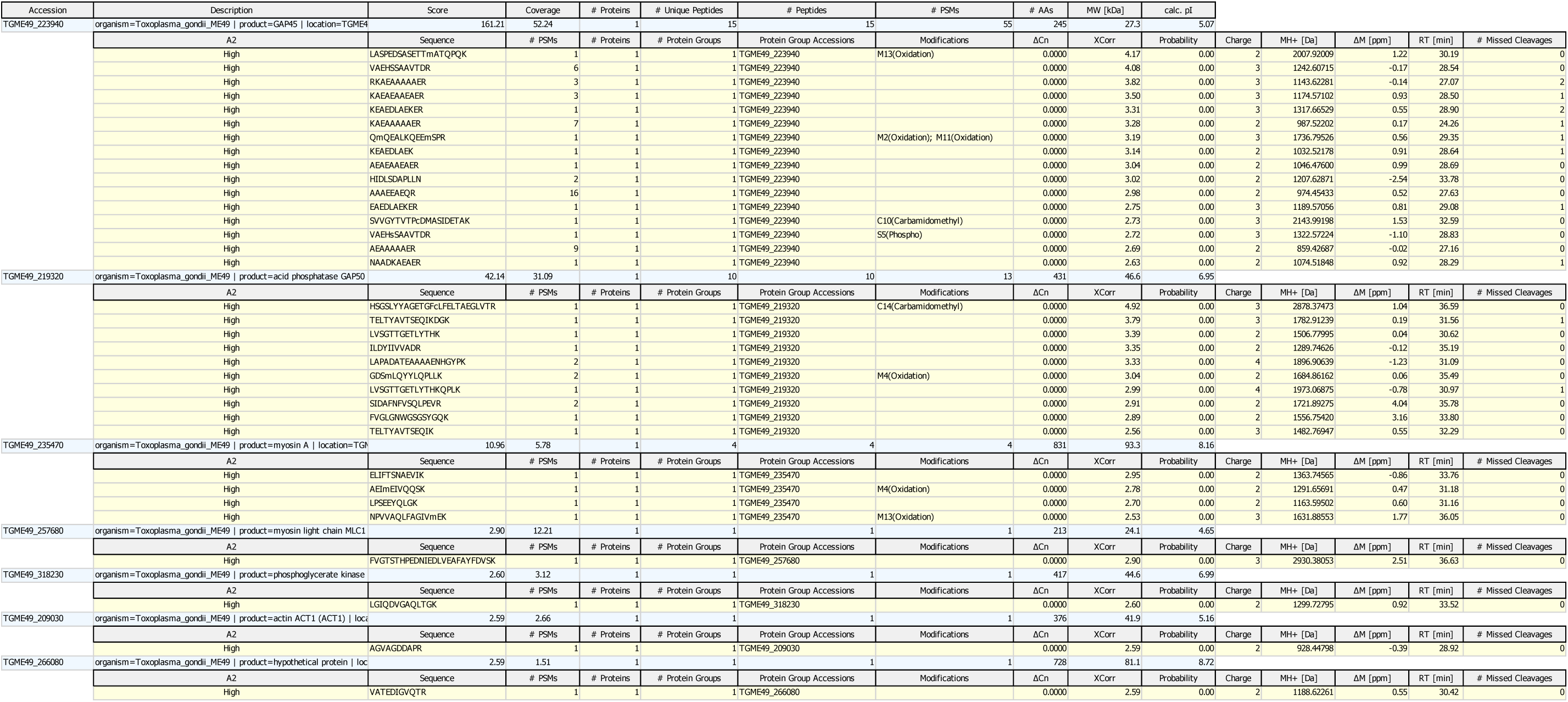
Identification of protein bands recovered in the anti-FLAG immunoprecipitates from WT and C(8,11)S parasites. Protein identification data for the nine excised bands shown in Fig. S6 are summarized in Table S1, and the raw LC/MS-MS results for each individual band are provided in Tables S2-S10.

**Supplementary Tables 1-10:**
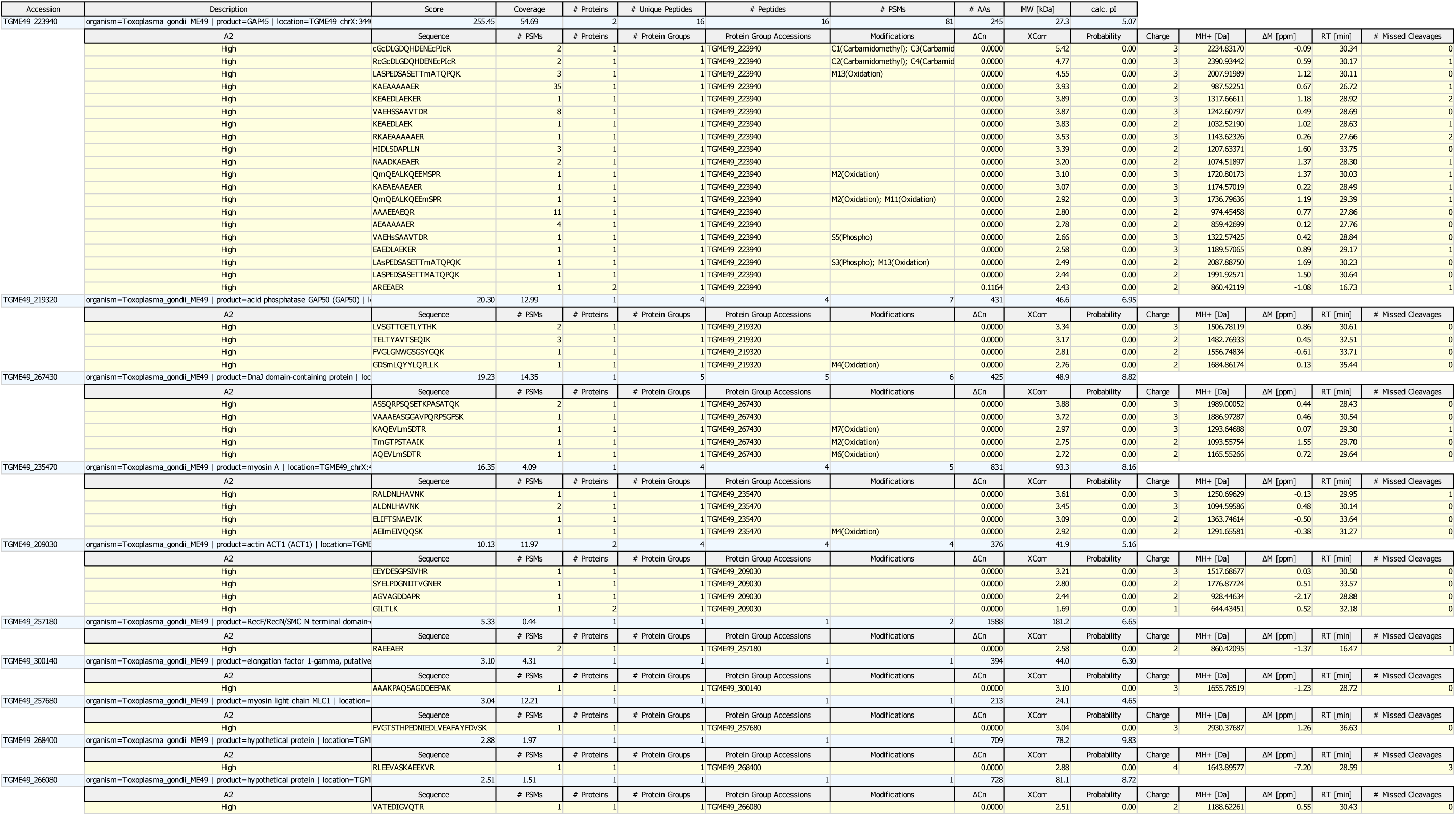
Identification of protein bands recovered in the anti-FLAG immunoprecipitates from WT and C(8,11)S parasites. Protein identification data for the nine excised bands shown in Fig. S6 are summarized in Table S1, and the raw LC/MS-MS results for each individual band are provided in Tables S2-S10.

**Supplementary Tables 1-10:**
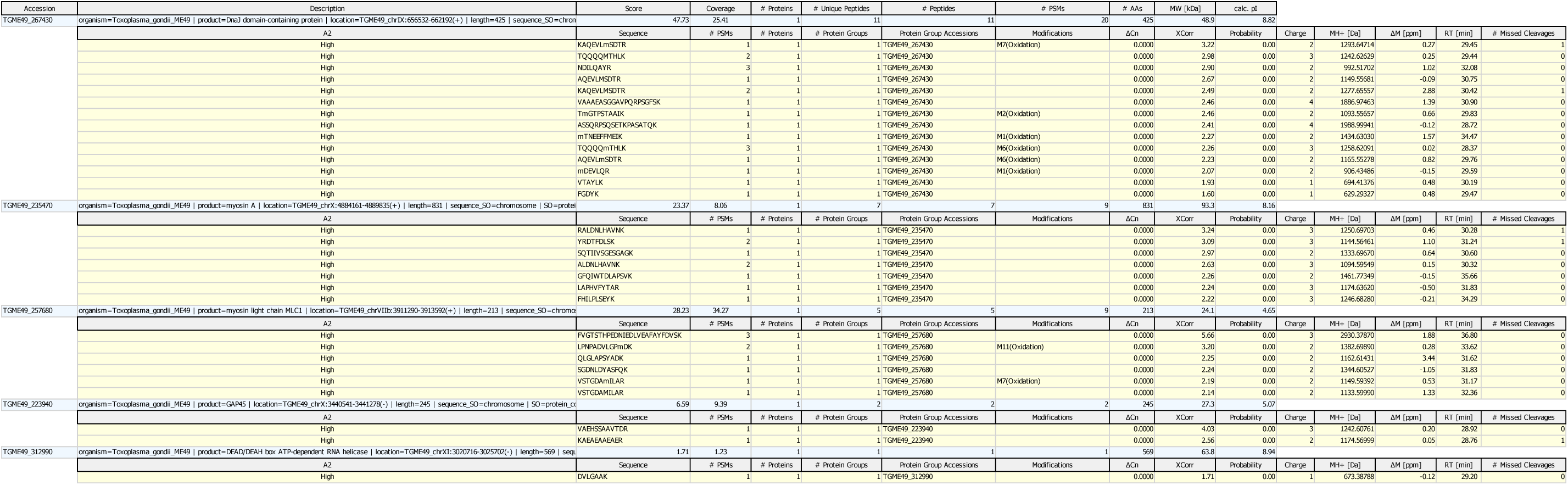
Identification of protein bands recovered in the anti-FLAG immunoprecipitates from WT and C(8,11)S parasites. Protein identification data for the nine excised bands shown in Fig. S6 are summarized in Table S1, and the raw LC/MS-MS results for each individual band are provided in Tables S2-S10.

**Supplementary Tables 1-10:**
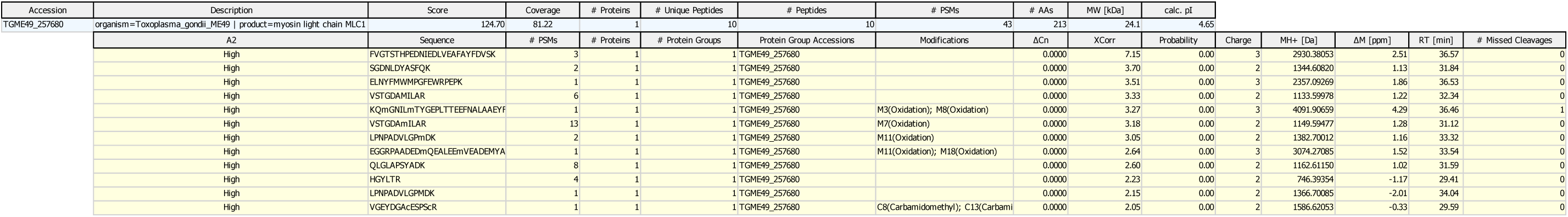
Identification of protein bands recovered in the anti-FLAG immunoprecipitates from WT and C(8,11)S parasites. Protein identification data for the nine excised bands shown in Fig. S6 are summarized in Table S1, and the raw LC/MS-MS results for each individual band are provided in Tables S2-S10.

**Supplementary Table 11.**
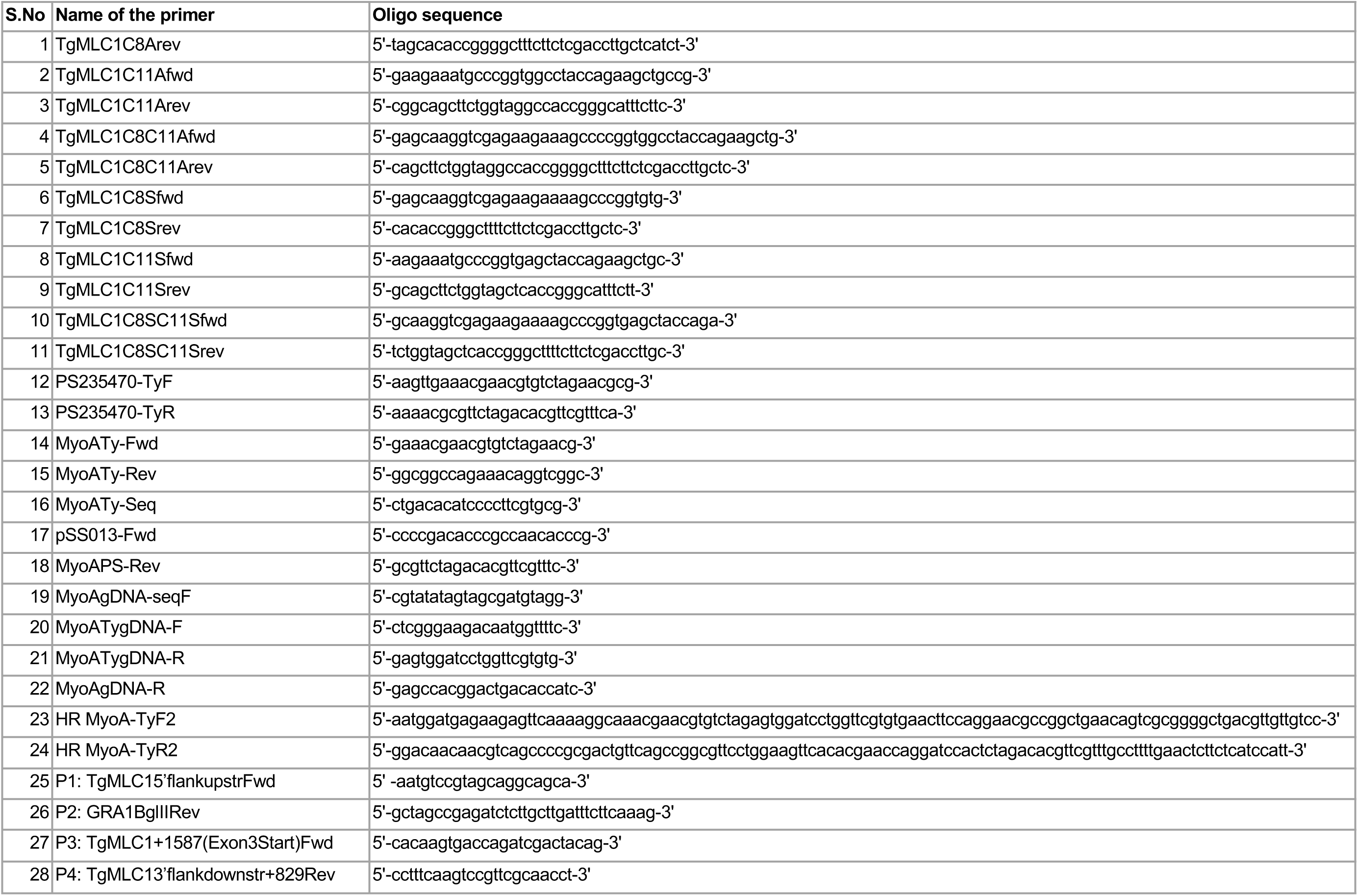
List of oligonucleotides used in this study.

## Acknowledgements

This work was supported by US Public Health Service grants AI139201 to GEW and GM111703 to MB. The Vermont Genetics Network Proteomics Facility is supported through NIH grant P20GM103449 from the INBRE Program of the National Institute of General Medical Sciences. The funders had no role in study design, data collection and interpretation, or the decision to submit the work for publication.

